# “Descending Enkephalinergic Neurons in Barrington’s Nucleus Gate Sex-Biased Control of Micturition”

**DOI:** 10.1101/2025.05.16.654570

**Authors:** Nataliya Klymko, Andrea M. Sartori, Mihoko Leon, Cassandra N. Seifert, Richard Lee, John C. Mathai, Anne M.J. Verstegen

**Affiliations:** Division of Nephrology, Department of Medicine, Beth Israel Deaconess Medical Center and Harvard Medical School, Boston, MA 02215, USA; University of Maryland School of Medicine, Baltimore, MD 21201, USA

## Abstract

Barrington’s nucleus (Bar) is a brainstem hub essential for lower urinary tract (LUT) function, yet the molecular identity and functional specialization of its neuronal subtypes remain poorly defined. Here, we construct a single-nucleus transcriptional atlas of Bar and identify an excitatory *Penk-* expressing population (Bar-*Penk*) critical for LUT regulation. Bar-*Penk* neurons are selectively active during voiding, when the external urethral sphincter (EUS) relaxes, and their optogenetic activation elicits time-locked suppression of EUS activity. Chemogenetic activation of these neurons induces a sex-specific, aberrant pattern of micturition, whereas targeted ablation impairs voluntary marking behavior in response to female cues. We observe multiple regions involved in visceromotor regulation and behavioral state control that project to Bar-*Penk* neurons, supporting their role as integrators of internal state and environmental context that drive urinary output. These findings provide new insights into the brainstem circuits that shape reflexive and voluntary micturition and highlight how discrete neuronal subtypes contribute to sexually dimorphic regulation of LUT function.

## Introduction

Few bodily functions are as tightly choreographed by brain-body interplay as urination which depends on continuous communication between the lower urinary tract (LUT) and the brain [1]. Barrington’s nucleus (Bar), located in the pontine tegmentum, innervates several visceral organs, including the bladder, urethra, urethral sphincters, colon, and reproductive organs [2–5]. The LUT relies on neural input from Bar to coordinate the muscles that control bladder function, which enables the socially appropriate and safe elimination of urine [6, 7]. When Bar is bilaterally lesioned, cats and rodents retain urine indefinitely [8, 9]. In humans, the LUT’s reliance on supraspinal control is evident from its vulnerability to neurogenic injuries, including stroke, neurodegenerative disorders, spinal cord injury, and aging [10–14]. However, despite the high prevalence and quality-of-life impact of LUT disorders, the neural control of LUT function remains underexplored.

On the ascending limb of the micturition reflex pathway, sensory information from bladder afferents is relayed via spinal interneurons in the lumbosacral cord to the midbrain periaqueductal gray (PAG), where it is further integrated. PAG neurons, in turn, project directly to Bar [15, 16]. In the descending pathway, Bar neurons send projections to spinal preganglionic motor neurons, which ultimately innervate the bladder, and interneurons that control somatic external urethral sphincter (EUS) motor neurons [17–21]. Upon the transition from storage to voiding, activation of Bar neurons drives detrusor contraction while simultaneously relaxing the urethral sphincters, facilitating urine release [19, 20]. The micturition reflex is under voluntary control, allowing humans and rodents, among other animals, to regulate the timing of urination based on context. This voluntary regulation is further exemplified by rodents’ use of urine marking for social communication [22–25].

Bar is predominantly glutamatergic [26]. Bar*^Vglut2^* neurons drive micturition, and their loss leads to severe urinary retention [27]. Neurons expressing *corticotropin-releasing hormone* (*Crh*) have long served as a proxy for Bar neurons overall [26, 28, 29], and their activation can induce both voiding and non-voiding bladder contractions depending on the phase of the micturition cycle [24, 25, 27, 30]. *Estrogen receptor alpha* (*Esr1*) is also abundantly expressed in Bar [31]; optogenetic stimulation of Bar*^Esr1^* neurons triggers EUS bursting and promotes urination, whereas their inhibition prevents EUS relaxation and interrupts ongoing voiding [25]. Recently, it was shown that distinct subpopulations of Bar*^Esr1^* exist, with respective projections via the pelvic nerve to control bladder-urethra activity and via the pudendal nerve to regulate EUS function [32].

Despite decades of research, the neuronal composition of Bar remains elusive, limiting our understanding of the functional roles of specific groups within this key brainstem nucleus for LUT control. Here, we characterize a distinct population of glutamatergic Bar neurons marked by *proenkephalin (Penk,* Bar*^Penk^*) expression. These neurons exhibit a sexually dimorphic influence on urinary behaviors, selectively promote EUS relaxation, and are essential for voluntary scent-marking in male mice. Our findings establish Bar*^Penk^* neurons as a specialized functional entity within Bar, revealing the neural mechanisms underlying both reflexive and voluntary micturition.

## Results

### Resolving neuronal diversity in Barrington’s nucleus through single-nucleus transcriptional profiling

Despite its critical role in LUT regulation, a comprehensive molecular atlas of Bar is lacking. Recent transcriptomic efforts, such as the atlas of the murine pontine tegmentum by Nardone et al. (2024) [33], include Bar within the broader brainstem region. However, they do not resolve its cellular heterogeneity, likely due to the nucleus small size relative to the large area captured. To identify the neuronal populations within Bar potentially involved in LUT control, we analyzed a Bar-enriched subset of the published single-nucleus RNA sequencing (snRNA-seq) dataset (GEO <u>GSE226809</u>) [33]. After pre-processing and quality controls, we performed preliminary clustering, removed clusters containing low-quality cells or doublets, and obtained a final dataset comprising 60,135 nuclei. Our analysis identified 39 clusters representing nine major cell types based on the expression of known marker genes (**Fig. 1a**). These included: ependymal cells (*Cfap43, Dnah12, Ccdc153*), fibroblasts (*Dcn, Cped1, Col3a1*), microglia (*Ctss, P2ry12, Csf1r*), endothelial cells (*Flt1, Slco1c1, Ly6c1*), astrocytes (*Ntsr2, Slco1c1, Ly6c1*), oligodendrocyte precursor cells (OPCs, *Pdgfra, Cspg4, Tnr*), oligodendrocyte progenitors (Oligo Prog, *Ust, Enpp6, Mag*), oligodendrocytes (*Mag, Plp1, Mog*), and neurons (*Snap25, Map2, Syt1*) (**Fig. 1b**).

**Figure 1.**
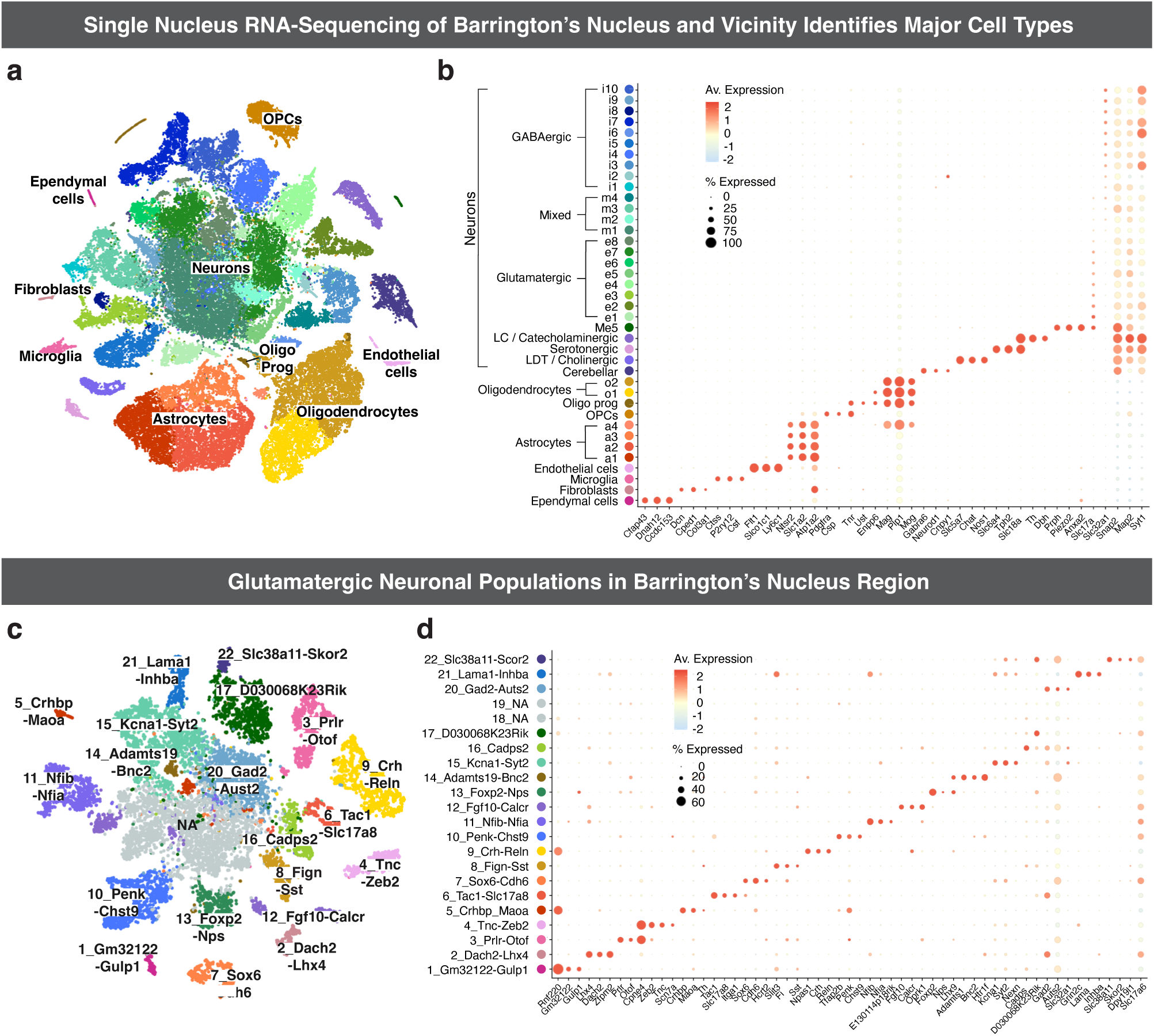
Resolving neuronal diversity in Barrington’s nucleus through single-nucleus transcriptional profiling. **a.** t-SNE plot of 60,135 nuclei from the Bar region. Each of the 39 clusters represents a distinct population of cells based on transcriptional profiles, grouped into 9 major cell types. **b.** Dot plot of canonical marker genes for major cell types. **c.** t-SNE plot displaying sub-clustering of glutamatergic neurons from (**a**), comprising 13,076 nuclei. **d.** Dot plot showing the top three most unique and/or highly expressed marker genes for each glutamatergic cluster. In (**b**, **d**), the color intensity represents average gene expression per cluster; the dot diameter reflects the percentage of nuclei expressing the gene. **Abbreviations:** LC, locus coeruleus; LDT, laterodorsal tegmental nucleus; Me5, mesencephalic nucleus of the trigeminal nerve; Oligo Prog, Oligodendrocyte progenitors; OPCs, oligodendrocyte precursor cells; sn-RNA-seq, single-nucleus RNA sequencing. *See also Supplementary Fig. 1 and Supplementary Table 1. Source data are provided as a Source Data file*.

The 27 neuronal clusters were further classified into eight subgroups based on gene expression profiles and anatomical location: GABAergic (*Vgat; Slc32a1)*; glutamatergic neurons (*Vglut2; Slc17a6*); mixed clusters (containing both *Vgat+* and *Vglut2+* cells); cerebellar neurons (*Gabra6, Neurod1, Cnpy1)*; laterodorsal tegmentum cholinergic neurons (*Slc5a7, Chat, Nos1*); a serotonergic population likely originating from the raphe nuclei rostral to Bar (*Slc6a4, Tph2, Slc18a2*); catecholaminergic neurons, associated with the locus coeruleus (LC; *Slc18a2, Th, Dbh*); and Me5 neurons, characterized by expression of *Prph, Piezo2*, and *Anxa2* (**Fig. 1b**).

### Identification of glutamatergic neuronal subtypes in Bar

To reveal the excitatory neuronal subpopulations of Bar that may contribute to LUT control, we reclustered neurons from clusters in which ≥10% of cells expressed the glutamatergic marker *Slc17a6* (*Vglut2*). We excluded non-neuronal cells, GABAergic clusters, and populations identified by markers of neighboring structures, including LC, cerebellar, cholinergic, and Me5 neurons. Mixed clusters co-expressing *Vgat* and *Vglut2* were retained to capture potential hybrid populations. Clustering of the resulting dataset (13,076 nuclei) yielded 22 distinct neuronal subgroups (**Fig. 1c, d; Supplementary Fig. 1a, b).** The neuronal populations were then mapped to their spatial location using RNAscope *in situ* hybridization (ISH), and gene expression patterns were cross-referenced with the Allen Mouse Brain Atlas [34] to identify genes that could be used for targeting the different neuron groups (**Supplementary Table 1**). This approach revealed six glutamatergic neuronal populations residing in or near Bar: “3_Prlr-Otof”, “6_Tac1-Slc17a8”, “9_Crh-Reln”, “10_Penk-Chst9”, “12_Fgf10-Calcr”, and “13_Foxp2-Nps”.

While the analysis identified the well-characterized *Crh*-expressing population in Bar [27, 30, 33], it did not detect a distinct *Esr1*-expressing cluster; rather, *Esr1* was detected across multiple Bar clusters, with highest expression detected in cluster “9_Crh-Reln” (**Supplementary Fig. 1c**). In contrast to previous reports [25], we observed high levels of *Esr1* expression in and around Bar, with substantial overlap between *Crh* and *Esr1*-positive neurons (**Supplementary Fig. 1d, e**; *88%, n = 3*). Extensive labeling in the *Esr1-Cre* mouse line following injection of a Cre-dependent AAV into Bar (**Supplementary Fig. 1f;** n = 2) further confirmed abundant *Esr1* expression in the region.

### Spinally projecting neuronal populations in Barrington’s nucleus

Identification of the neuronal populations within and immediately adjacent to Bar lays the groundwork for investigating their potential roles in LUT regulation. To this end, we traced descending projections from each population to spinal regions critical for bladder and EUS control, specifically, the intermediolateral cell column (IML) and dorsal gray commissure (DGC) in the lumbosacral spinal cord, where motor neurons and interneurons innervating the LUT are located [3, 4] (**Fig. 2a, b**). First, we confirmed that glutamatergic, but not GABAergic, neurons in Bar send descending projections to the spinal cord. Robust axonal labeling from *Vglut2+* pan-Bar neurons was observed in the L6-S3 spinal segments (**Supplementary Fig. 2c**), whereas no axons from *Vgat+* neurons were detected in the lumbosacral spinal cord (**Supplementary Fig. 2d**). Thus, glutamatergic Bar neurons provide the primary descending input to spinal circuits controlling LUT function, while adjacent inhibitory populations [33] do not contribute direct projections.

**Figure 2.**
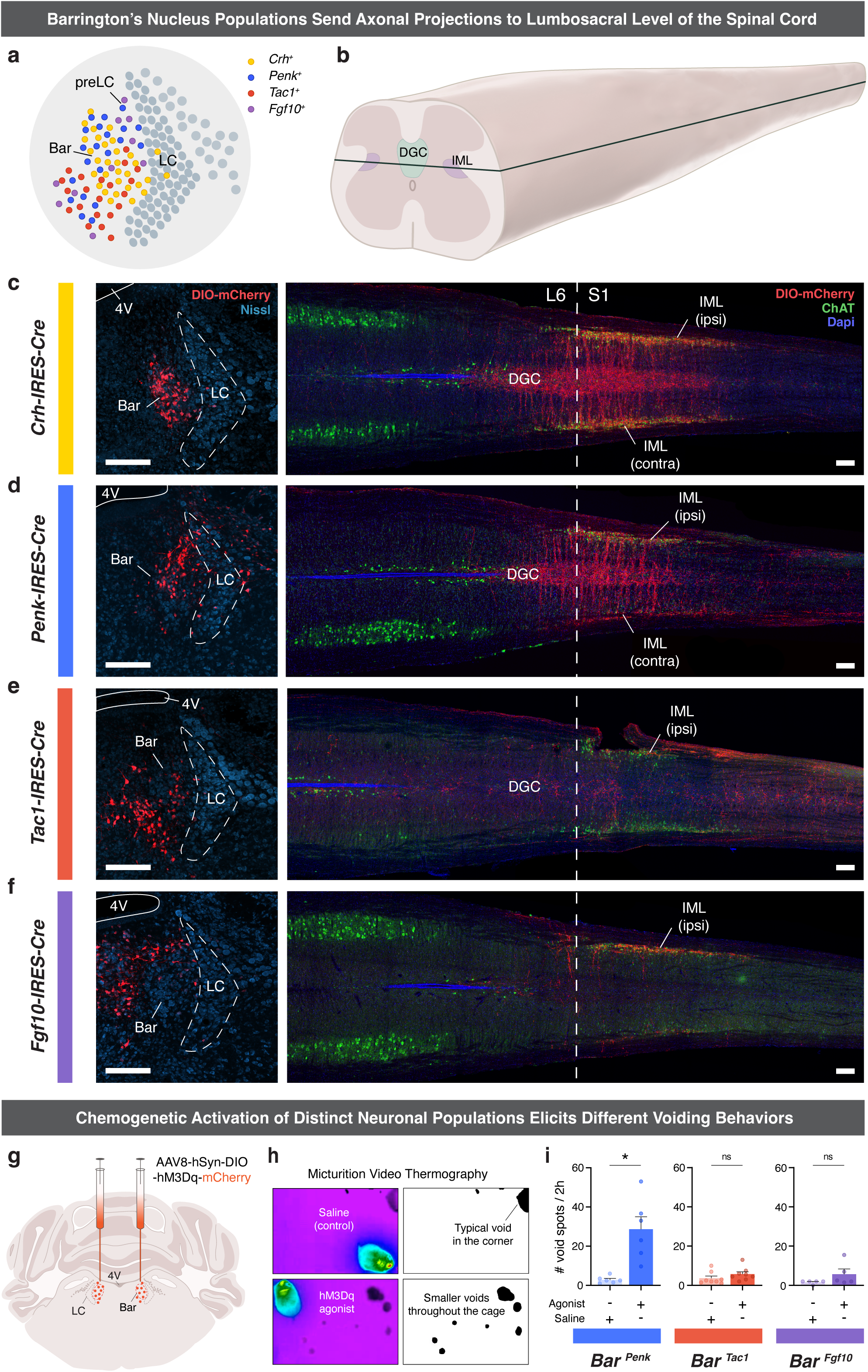
Spinally projecting neuronal populations of Bar and their role in micturition control. **a.** Schematic of Bar region showing spinally projecting glutamatergic populations based on marker gene expression. **b.** Illustration of the spinal cord displaying a transverse section (black line) through DGC and IML at the lumbosacral level. **c-f.** DIO-mCherry labeling (red) in Bar of Cre-driver mice, representing neuronal populations (left); neuronal cell bodies are counterstained with Nissl (teal blue). Corresponding axonal projections to lumbosacral levels of the spinal cord (right); cholinergic neurons stained for ChAT (green) and nuclei for DAPI (blue). Ipsilateral (ipsi) and contralateral (contra) projections are indicated; dashed line marks L6/S1 spinal cord level (**c**; Bar*^Crh^*, *n = 3*; **d**; Bar*^Penk^*, *n = 8*; **e**; Bar*^Tac1^*, *n = 3*; **f**; Bar*^Fgf10^*, *n = 4*). **g**. Schematic of bilateral DIO-hM3Dq injections into Bar. **h.** Micturition Video Thermography (MVT) examples, with thresholded images of void spots displayed on the right. **i.** Averaged micturition frequency for Bar*^Penk^* (*n = 6, *p = 0.031*), Bar*^Tac1^*(*n = 8, p = 0.22*), and Bar*^Fgf10^* (*n = 5, p = 0.38*) from 2h MVT trials with saline or hM3Dq agonist; *mean ± SEM, two-sided Wilcoxon matched-pairs signed rank test across all panels.* Source data are provided as a Source Data file. **Scale bars**: 200 μm. **Abbreviations**: 4V, 4th ventricle; Bar, Barrington’s nucleus; ChAT, choline acetyltransferase; DGC, dorsal gray commissure; IML, intermediolateral column; LC, locus coeruleus; preLC, pre-locus coeruleus. *See also Supplementary Fig. 2, Supplementary Table 2*.

Using the *Crh* marker to target the ‘*Crh-Reln*’ population, we found *Crh*-expressing neurons to be distributed throughout the entire rostral-caudal extent of Bar. Axons from Bar*^Crh^* neurons descend ipsilaterally, heavily innervating the ipsilateral sacral IML; there, the fibers cross over to the DGC and extend to the contralateral IML at spinal segments L6, S1, and S2. **(Fig. 2c).** We then utilized the *Penk-IRES2-Cre* and *Tac1-IRES2-Cre* mouse lines to target cells of the ‘*Penk-Chst9’* and *‘Tac1-Slc17a8’* populations (**Fig. 2d, e**). *Penk+* neurons were predominantly localized in the dorsomedial aspect of Bar, with additional presence along the ventromedial border of the Nissl-defined oval Bar nucleus (**Fig. 2a**). Descending projections from the Bar*^Penk^* neurons were concentrated at spinal levels L6-S2, where they reached segments with ChAT+ bladder motor neurons in the ipsilateral and contralateral IML and DGC (**Fig. 2d**). *Tac1*+ reporter-labeled neurons were located in the ventral portion of Bar and along its ventromedial border (**Fig. 2a**). Projections from these neurons were also observed, but appeared more diffuse and less anatomically confined, with axons extending rostrally and caudally, reaching as far as S3/S4 spinal segments (**Fig. 2e**).

To target the ‘*Fgf10-Calcr*’ population, we used the *Fgf10-IRES-Cre* mouse line (**Supplementary Fig. 2a**). *Fgf10*+ neurons were distributed in the dorsal aspect of Bar and in the pre-LC (**Fig. 2a**) [35]. Sparse axonal projections reached the sacral IML, likely reflecting the limited number of *Fgf10*+ cells within the core of Bar (**Fig. 2f**). Finally, *Foxp2+* neurons, corresponding to the ‘Foxp2-Nps’ population, were labeled dorsolateral to Bar in the pre-LC [26, 36–38], and at the caudal extent of Bar. These *Foxp2*+ neurons did not project to the lumbosacral spinal cord, suggesting they do not exert direct control over the LUT motor neurons (**Supplementary Fig. 2e**).

While we confirmed using the *Prlr-P2A-Cre* mouse line that *Prlr*-expressing neurons project to the spinal cord (**Supplementary Fig. 2b, f**), further analysis of GFP expression in *Prlr-P2A-Cre::L10-GFP* mice revealed a large number of labeled neurons distributed throughout the pontine tegmentum (**Supplementary Fig. 2h**). This coincides with the high number of Cre-expressing neurons observed following DIO-mCherry injections in this transgenic line (**Supplementary Fig. 2f**). Additionally, we found a high degree of colocalization of *Crh* and *Penk* mRNA expression with Prlr-GFP neurons in Bar (**Supplementary Fig. 2i**). These results suggest that expression of *Prlr* is not exclusive to a single cell type or confined to the Bar nucleus, but rather that prolactin receptors are present across multiple subpopulations, with varying levels of expression. Consistent with *Esr1* marking a broad Bar population, *Esr1-Cre* mice showed robust reporter labeling in Bar and dense axonal projections to both the IML and DGC, comparable to that observed in *Vglut2-IRES-Cre* mice (**Supplementary Fig. 2c, g**).

### Functional screening of Bar populations for their role in micturition behavior

To evaluate the role of spinally projecting Bar subpopulations in LUT control, we selectively activated different neuronal populations using excitatory DREADDs (Designer Receptors Exclusively Activated by Designer Drugs; hM3Dq), delivered bilaterally into Bar of the respective Cre-driver mice (**Fig. 2g, h**). The Bar*^Crh^* population was not included, as we previously showed its activation with DREADDs to markedly increase voiding frequency (from 2.3 to 8.8 voids; n = 6) [27]. Activation of Bar*^Penk^* neurons resulted in aberrant voiding behavior, with a striking increase in the number of void spots (**Fig. 2i**, left panel). No significant changes in voiding frequency were observed when Bar*^Tac1^* and Bar*^Fgf10^* populations were chemogenetically stimulated (**Fig. 2i**, middle and right panels). Lastly, stimulation of *Prlr+* neurons in Bar produced a strong increase in micturition frequency, as would be expected when targeting multiple functional populations in Bar (**Supplementary Fig. 2j**).

### *Penk* expression defines a distinct neuronal population in Bar

Given the distinct behavioral effects observed with Bar*^Penk^* activation, we sought to further characterize this neuronal population. First, we examined their anatomical organization relative to the more extensively studied Bar*^Crh^*neurons (**Fig. 3a, b**). Along the anterior-posterior axis of Bar, *Crh+* neurons are distributed uniformly across the entire extent of the nucleus (Bregma −5.35 to −5.65 mm). In contrast, *Penk+* cells form a distinct cluster in the dorsomedial portion of the Nissl-defined Bar nucleus and are more concentrated at central and caudal levels, with peak density observed around Bregma −5.55 (**Fig. 3c**). To further delineate their molecular identity, we evaluated the overlap of the *Penk*-expressing neurons with known Bar markers using RNAscope ISH and immunohistochemistry (**Fig. 3d**). Virtually all Bar*^Penk^* neurons expressed *Vglut2* mRNA (95%, n = 6), confirming their excitatory nature, while only a small subset (5%) co-expressed *Crh* mRNA (n = 6). Immunohistochemistry revealed that just over half of Bar*^Penk^* neurons were positive for Esr1, another marker for glutamatergic Bar neurons, with no significant difference between males and females (**Supplementary Fig. 3a, b;** 57%, n = 4 males, 3 females). Overall, *Penk+* neurons comprise one-fifth of the total Bar population, while *Crh+* neurons account for 40%, and other populations (negative for both markers) make up the remaining 38%. Remarkably, only 1% of the total Bar neuronal pool expressed both *Penk* and *Crh* (**Fig. 3e**). *Esr1*+ neurons comprised 75% of the overall Bar population and showed substantial overlap with both *Penk* and *Crh* (**Supplementary Fig. 3c-e**), consistent with *Esr1* being broadly expressed in Bar (**Supplementary Fig. 1d-f, Supplementary Fig. 2g**).

**Figure 3.**
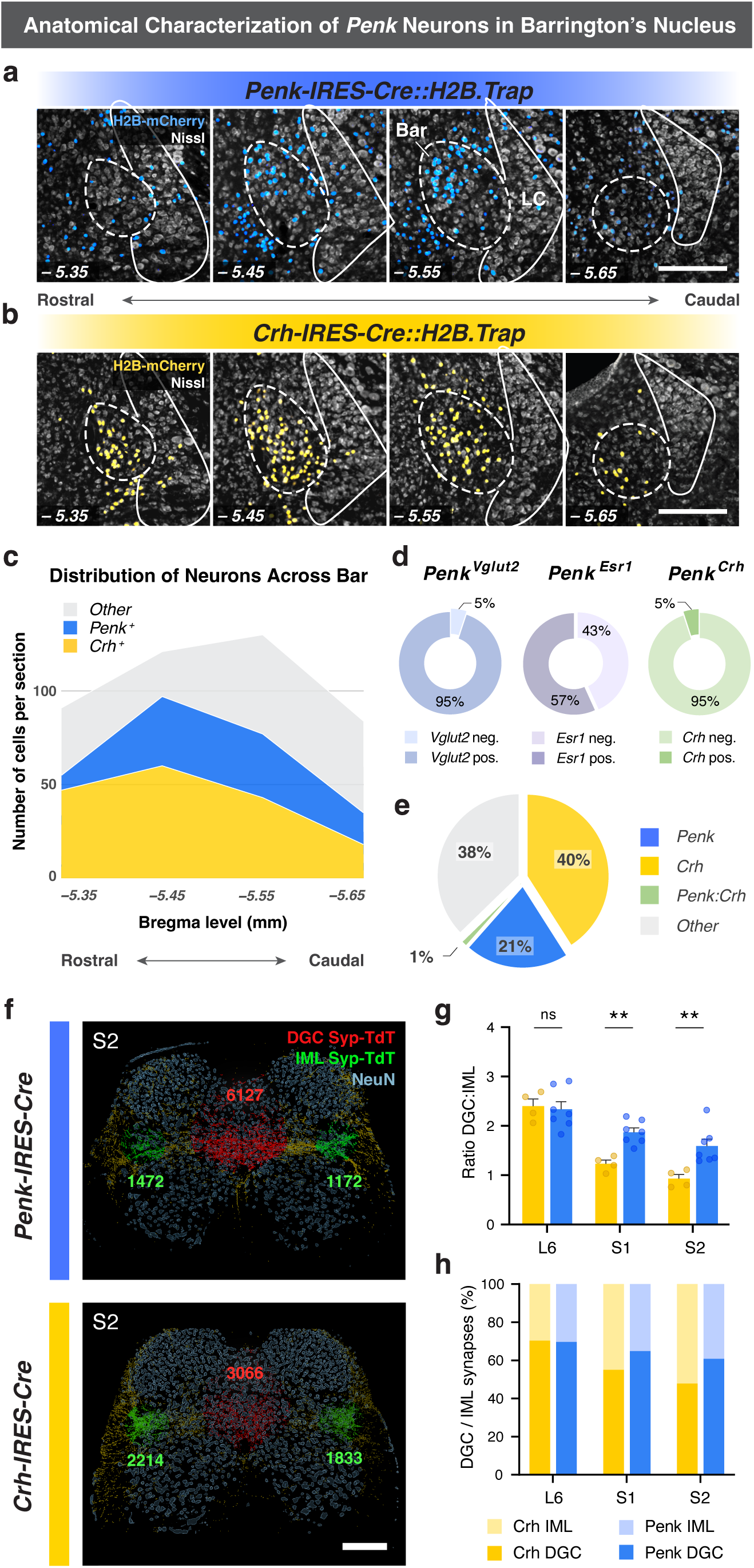
*Penk* expression defines a distinct neuronal population in Bar. **a-b.** Anatomical distribution of *Penk+* (blue, *n = 3*) and *Crh*+ (yellow, *n =3*) neurons across Bregma levels in Bar; neuronal cell bodies are counterstained with Nissl (gray). **c.** Quantification of Bar*^Penk^*(*n = 3*), Bar*^Crh^* (*n = 3*), and other (unlabeled) Bar neurons across four Bregma levels. **d.** Overlap of *Bar^Penk^*neurons with known Bar markers: *Vglut2* (*95 ± 0.8%, n = 6*), *Esr1* (*57 ± 3.4%, n = 7*), and *Crh* (*5 ± 0.7%, n = 6*). **e.** Proportion of *Penk+* and *Crh+* neurons in Bar. **f.** Imaris-processed images of the sacral spinal cord showing the distribution of Bar*^Penk^* (top) and Bar*^Crh^*(bottom) synaptic terminals within the DGC (red spots) and IML (green spots) following injection of AAV-DIO-mSyp1-tdTomato into Bar of *Penk-IRES2-Cre* and *Crh-IRES-Cre* mice. Numbers denote synaptic spot counts within the DGC and IML in the representative sections shown; spots outside the quantified ROIs appear in yellow. **g.** DGC:IML synapse ratio for Bar*^Penk^* (blue, *n = 7*) and Bar*^Crh^* (yellow, *n = 4*) across lumbosacral spinal levels (L6, S1, and S2). *Mean ± SEM; p = 0.98; **p = 0.0078; **p = 0.0058, two-way ANOVA with Sidak’s multiple-comparisons test.* **h.** Average distribution of synapses between the DGC and IML at each spinal level, expressed as the % of terminals within that level, for Bar*^Penk^* (blue) and Bar*^Crh^*(yellow) **Scale bars**: 200 μm. **Abbreviations**: Bar, Barrington’s nucleus; DGC, dorsal gray commissure; IML, intermediolateral column; LC, locus coeruleus. *See also Supplementary Fig. 3. Source data are provided as a Source Data file*.

To further characterize Bar*^Penk^* axonal innervation of the lumbosacral spinal cord, we compared the distribution of synaptic boutons from Bar*^Penk^* and Bar*^Crh^* neurons at L6-S2 spinal levels. Although both populations projected to the DGC and IML (**Fig. 2c, d**), their relative patterns of innervation diverged. At L6, where the IML starts to emerge, the DGC vs. IML distribution of synapses was similar between the two populations. However, at S1 and S2, the patterns differed significantly, with Bar*^Penk^* boutons enriched in the DGC and Bar*^Crh^* boutons showing greater enrichment in the IML (**Fig. 3f-h, Supplementary Fig. 3f, g**). Sparse bouton labeling was also observed in the dorsal horn, with Bar*^Crh^* boutons preferentially localized to the superficial laminae and Bar*^Penk^* boutons distributed more broadly across the dorsal horn. Thus, despite overlapping spinal targets, the two populations exhibit distinct subregional distributions of synaptic input.

### *Penk* neurons in Bar activate specifically during voiding

To determine whether the physiological, *in vivo* activity of Bar*^Penk^* neurons correlates with specific phases of the bladder fill-void cycle, we simultaneously monitored neural activity and bladder pressure using fiber photometry and cystometry in awake, freely behaving mice (**Fig. 4a-c**). Bar*^Penk^* neurons consistently activated around the time of void onset and remained active for the duration of the void (**Fig. 4d-f**). Notably, increases in neural activity did not precede rises in intravesical pressure; these findings suggest that Bar*^Penk^* neuron activity is closely correlated with voiding, potentially coordinating EUS relaxation rather than detrusor contractions. Small transient changes in GCaMP fluorescence also accompanied pressure changes during non-voiding contractions, but activity was attenuated compared to that observed during voids (**Supplementary Fig. 4a, b**).

**Figure 4.**
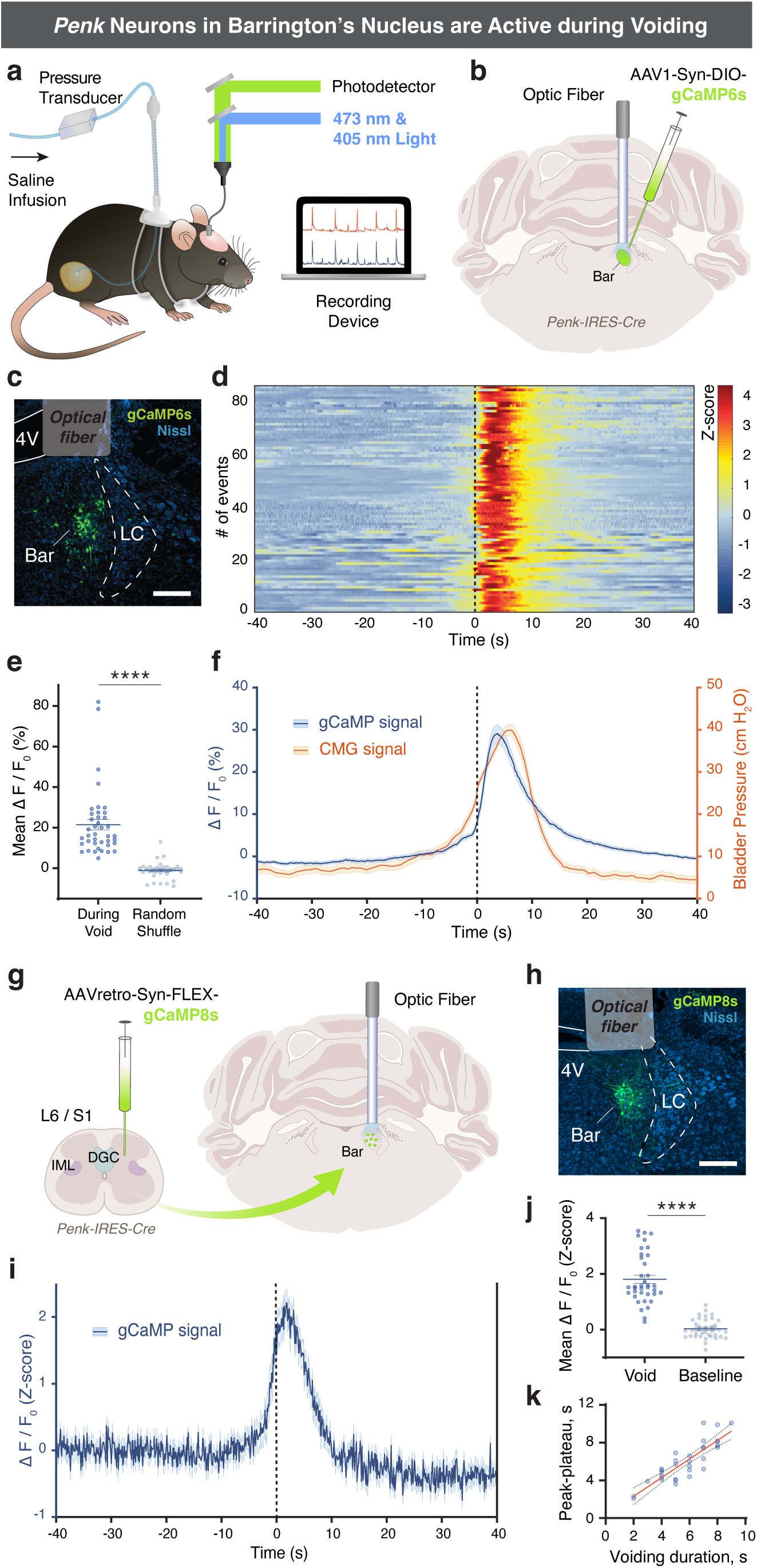
Bar*^Penk^* neurons selectively activate during voiding. **a.** Experimental setup combining fiber photometry and cystometry (CMG) to record neural activity= and bladder pressure simultaneously. **b.** Schematic showing unilateral injection of DIO-GCaMP6s= targeting Bar with optic fiber placement above. **c.** DIO-GCaMP6s expression in Bar of a *Penk-IRES-Cre* mouse, with the fiber positioned above. **d.** Heatmap of GCaMP6s signal (*ΔF/F_0_*,= converted to Z-score) before, during, and after individual voiding events. **e.** Average GCaMP6s signal (*ΔF/F_0_*) during voiding vs. at random times (*mean ± SEM*, *n = 40 events from 4 mice, equally weighted; ****p < 0.0001, two-sided Mann-Whitney test).* **f.** Averaged CMG trace (orange) and GCaMP6s signal (blue, *ΔF/F_0_*) across 86 voiding events from 4 mice (solid lines represent the mean, shaded areas - SEM). **g.** Experimental approach for retrograde targeting and fiber photometry of spinally projecting Bar*^Penk^* neurons: lumbosacral spinal injection of AAVretro-FLEX-GCaMP8s in *Penk-IRES-Cre* mice, followed by optic fiber implantation over Bar. **h.** GCaMP8s expression in retrogradely labeled Bar*^Penk^*neurons and fiber placement above. **i.** Averaged photometry trace (*z-scored ΔF/F_0_; mean ± SEM; n = 37 events from 6 mice*) from spinally projecting Bar*^Penk^*neurons recorded during MVT, aligned to void onset (t = 0, first thermal frame where urine spot appears). **j.** Mean *z-scored ΔF/F_0_* during voiding vs. pre-void baseline (*mean ± SEM; n = 37 events from 6 mice; ****p < 0.0001, two-sided Wilcoxon matched-pairs signed rank test*). **k.** Relationship between voiding duration and peak-plateau duration of Bar*^Penk^* activity (sustained high GCAMP signal; *n = 32 events, from 6 mice*). Red line indicates linear regression fit; dotted lines indicate the 95% confidence band (*Pearson correlation r = 0.86, 95% CI 0.73 - 0.93; R² = 0.74; p < 0.0001*). In **(d, f, i)** dashed line marks approximate void onset. In (**c, h**), neuronal cell bodies are counterstained with Nissl (blue). **Scale bars**: 200 μm. **Abbreviations**: 4V, 4^th^ ventricle; Bar, Barrington’s nucleus; LC, locus coeruleus; MVT, Micturition Video Thermography. *See also Supplementary Fig. 4. Source data are provided as a Source Data file*.

Fiber photometry recordings specifically from the spinally-projecting Bar*^Penk^* neurons (**Fig. 4g, h**), coupled with video thermography, revealed a similar activity pattern, with robust activation time-locked to the void onset (**Fig. 4i, j; Supplementary Fig. 4c**). Moreover, the duration of peak-plateau neural activity strongly correlated with void duration, indicating that Bar*^Penk^* activity scales with the length of urine release (**Fig. 4k; Supplementary Fig. 4d**).

### Activation of Bar*^Penk^* neurons results in a sex-dependent behavioral phenotype

Having established that Bar*^Penk^* neurons project to the lumbosacral spinal cord and are active during voiding, we next asked whether their selective activation alters micturition behavior in male and female mice (**Fig. 5a, b**). In males, chemogenetic stimulation led to increased voiding frequency in some animals, although the average number of regular corner voids did not change (**Fig. 5c**). Urine volume per void was significantly reduced, which could suggest that prolonged Bar*^Penk^* activation impairs normal bladder emptying (**Fig. 5d**). Additionally, throughout the session, we observed numerous small urine spots consistent with leaks. These events occurred outside the typical voiding sequence with the spots scattered throughout the behavioral arena, suggestive of an incontinence-like phenotype. (**Fig. 5b, e, i**). In contrast, stimulation of the Bar*^Penk^* neurons in female mice produced no significant change in voiding frequency, urine volume per void, or leak occurrence between agonist and saline conditions (**Fig. 5f-h**). To control for potential off-target effects of the DREADDs agonist, we tested whether administration of the agonist in mice lacking viral expression influenced LUT function. No significant differences were observed between agonist-treated naïve animals and saline controls, in either males (n = 4 vs. 6) or females (n = 6 vs. 7) for any of the parameters tested (**Fig. 5c-h**).

**Figure 5.**
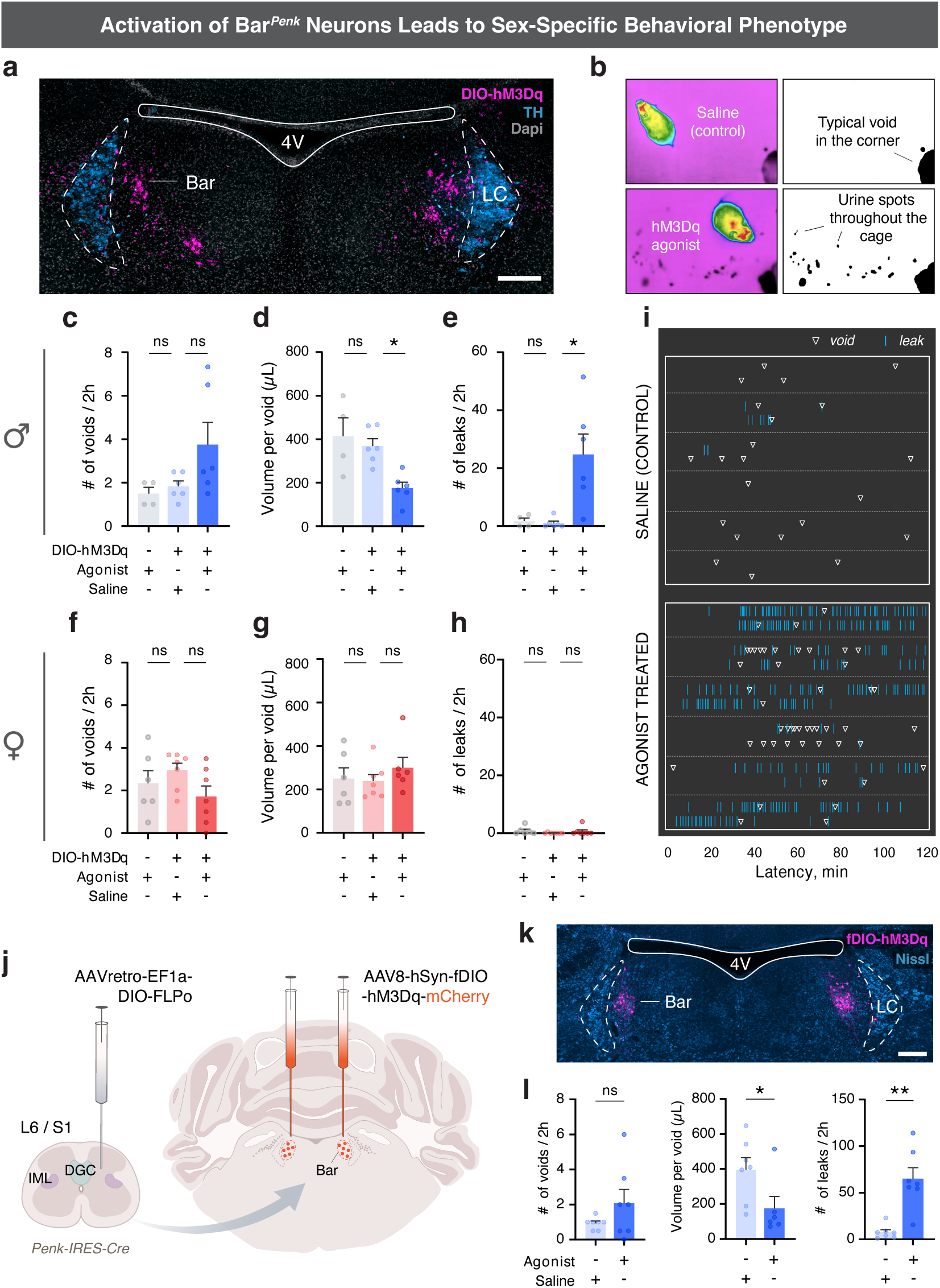
Activation of Bar*^Penk^* neurons triggers a sex-dependent behavioral phenotype. **a.** Representative image showing bilateral expression of DIO-hM3Dq (magenta) in Bar of a *Penk-IRES-Cre* mouse (*n = 6*). TH-immunoreactive LC neurons are shown in teal blue; nuclei counterstained with DAPI (gray). **b.** Representative MVT screenshots showing micturition behavior under saline control (top) and after hM3Dq agonist administration (bottom). **c-h.** Quantification of micturition parameters, including voiding frequency **(c, f)**, urine volume per void **(d, g)**, and number of leaks **(e, h)** after Bar*^Penk^* activation. Data are shown for males (**c, d, e;** *n = 6, p = 0.33, *p = 0.027, *p = 0.017,* respectively) and females (**f, g, h;** *n = 7, p = 0.17, p = 0.67, p > 0.99,* respectively). Off-target effects of hM3Dq agonist were tested in control animals lacking viral expression (**c, d, e;** *n = 4 males, all p > 0.99*; **f, g, h;** n = 6 females, *p = 0.97, p > 0.99, p = 0.44,* respectively). *Mean ± SEM, Kruskal-Wallis, followed by Dunn’s multiple comparisons tests across all panels.* **i.** Raster plot showing voiding behavior during 2h MVT runs under saline control (top) and agonist-treated (bottom) conditions (*n = 6*; corresponding quantification is shown in **c** and **e**). Each row represents one recording session (two per animal for each condition). Voids are shown as open white triangles and leaks as cyan ticks. **j.** Experimental approach for selective targeting and chemogenetic activation of spinally projecting Bar*^Penk^*neurons: bilateral AAVretro-DIO-FLPo injection into the lumbosacral spinal cord (one side shown) of *Penk-IRES-Cre* mice, followed 3 weeks later by bilateral injection of AAV8-fDIO-hM3Dq-mCherry into Bar. **k.** Flp-dependent hM3Dq-mCherry expression in spinally projecting Bar*^Penk^*neurons. **l.** Voiding frequency (left), volume per void (middle), and number of leaks (right) following activation of spinally projecting Bar*^Penk^*neurons (*mean ± SEM; n = 7; p = 0.51, *p = 0.035, **p = 0.0012, respectively; two-sided Mann–Whitney tests*). **Scale bars**: 200 μm. **Abbreviations**: 4V, 4th ventricle; Bar, Barrington’s nucleus; LC, locus coeruleus; MVT, Micturition Video Thermography; TH, tyrosine hydroxylase. *See also Supplementary Fig. 5. Source data are provided as a Source Data file*.

Selective activation of spinally projecting Bar*^Penk^*neurons in males (**Fig. 5j, k**) recapitulated the main effects observed after overall Bar*^Penk^* stimulation, reducing urine volume per void and increasing leak occurrence without significantly altering void frequency (**Fig. 5l**), indicating that activating the spinally projecting population is sufficient to produce the incontinence-like phenotype.

### Bar*^Penk^* neurons are essential for voluntary scent-marking behavior

To assess whether *Penk* neurons in Bar are necessary for normal LUT function, we ablated them using diphtheria toxin (dtA) (**Fig. 6a, b**). Conditional ablation of the Bar*^Penk^* neurons did not affect voiding frequency or void sizes in either sex (**Fig. 6c-f; Supplementary Fig. 6a-d**), in contrast to ablation of the entire glutamatergic Bar population, which leads to severe urinary retention with overflow incontinence [27].

**Figure 6.**
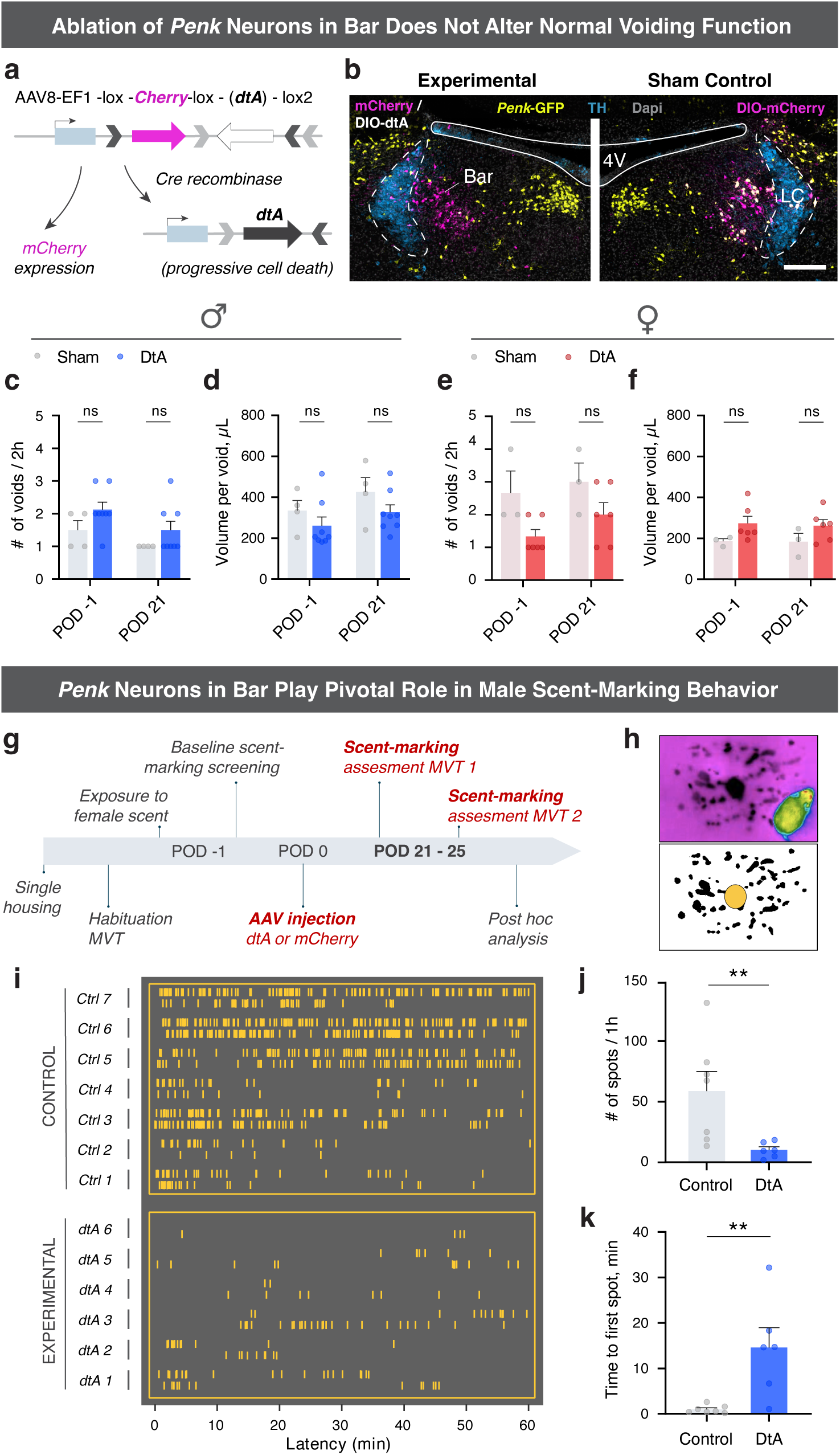
Bar*^Penk^* neurons are essential for voluntary scent-marking behavior. **a.** The viral construct of Cre-dependent diphtheria toxin A (dtA). **b.** Injection site images showing mCherry expression in Cre-negative cells and absence of dtA-infected Penk-GFP neurons (lime-green) in Bar (experimental, left). In sham controls (right), *Penk+* neurons co-express GFP and DIO-mCherry. One side is shown; both groups received bilateral injections. **c-f.** Average voiding frequency (**c, e**) and urine volume per void (**d, f**) before and after dtA-mediated Bar*^Penk^*ablation in males (**c, d;** *n = 8 vs. n 4;* in (**c**), *p = 0.34* (POD -1), *p = 0.42* (POD21); in (**d**), *p = 0.60* (POD - 1), *p = 0.60* (POD21)) and females (**e, f**; *n = 6 vs. n = 3*; in (**e**), *p = 0.22* (POD -1), *p = 0.30* (POD21); in (**f**), *p = 0.18* (POD -1), *p = 0.26* (POD21)). *Mean ± SEM*, *two-sided Mann-Whitney tests with Holm-Šídák correction for multiple comparisons across all panels.* **g.** Experimental timeline for investigating effects of Bar*^Penk^* ablation on scent-marking behavior. **h**. Natural scent-marking behavior of dominant male mice exposed to female mouse urine cues. **i.** Raster plot of marking events (gold lines) recorded during two 1h MVT trials at POD 21-25, comparing Bar*^Penk^*-ablated experimental animals to sham controls. **j, k.** Quantification of urine marks (**j**) and latency to the first mark (**k**) at POD 21-25 in experimental vs. control groups (*mean ± SEM*, *n = 6 vs. n = 7; **p = 0.0047, **p = 0.0058, two-sided Mann-Whitney tests*). **Abbreviations**: 4V, 4th ventricle; Bar, Barrington’s nucleus; dtA, diphtheria toxin A; LC, locus coeruleus; MVT, Micturition Video Thermography; POD, postoperative day; TH, tyrosine hydroxylase. *See also Supplementary Fig. 6. Source data are provided as a Source Data file*.

Since Bar*^Penk^* neuron activation led to the appearance of scattered urinary spots throughout the cage (**Fig. 5b, e**), and ablation did not disrupt normal voiding, we asked whether these neurons may play a role in voluntary marking behaviors. Scent-marking relies on rapid, inhibitory control over the striated EUS muscle, and male mice engage in prolific scent-marking to attract female mates [22, 23, 25]. To evaluate the role of Bar*^Penk^* neurons in scent-marking, we recorded marking behavior before and three weeks after bilateral injection of dtA or mCherry (**Fig. 6g, h; Supplementary Fig. 6e, f**). Ablation of the *Penk*-expressing neurons resulted in a significant reduction in the number of urine marks in response to a stimulus (female mouse urine), along with a pronounced delay in the initiation of marking (**Fig. 6i-k**), with the effects observed only following complete bilateral ablation (**Supplementary Fig. 6g-h**). These findings demonstrate that Bar*^Penk^* neurons are essential for this voluntary behavior in male mice.

### Photostimulation of Bar*^Penk^* neurons promotes relaxation of the external urethral sphincter

To elucidate whether activation of Bar*^Penk^* neurons affects EUS behavior, we performed electromyography (EMG) recordings of the EUS and cystometry (CMG) in awake, freely behaving mice, coupled with optogenetic stimulation (**Fig. 7a, b**). Photostimulation at 20Hz frequency produced a time-locked reduction in EUS activity on EMG compared to pre-stimulation baseline and recovery periods (**Fig. 7c, d; Supplementary Fig. 7a**), while intravesical pressure remained unchanged (**Fig. 7e**). Notably, 75% of stimulations resulted in EUS relaxation, whereas an increase in bladder pressure was observed in only 6% of trials (**Fig. 7f**). These results suggest that Bar*^Penk^* neurons are sufficient to induce sphincter relaxation independent of bladder contractions.

**Figure 7.**
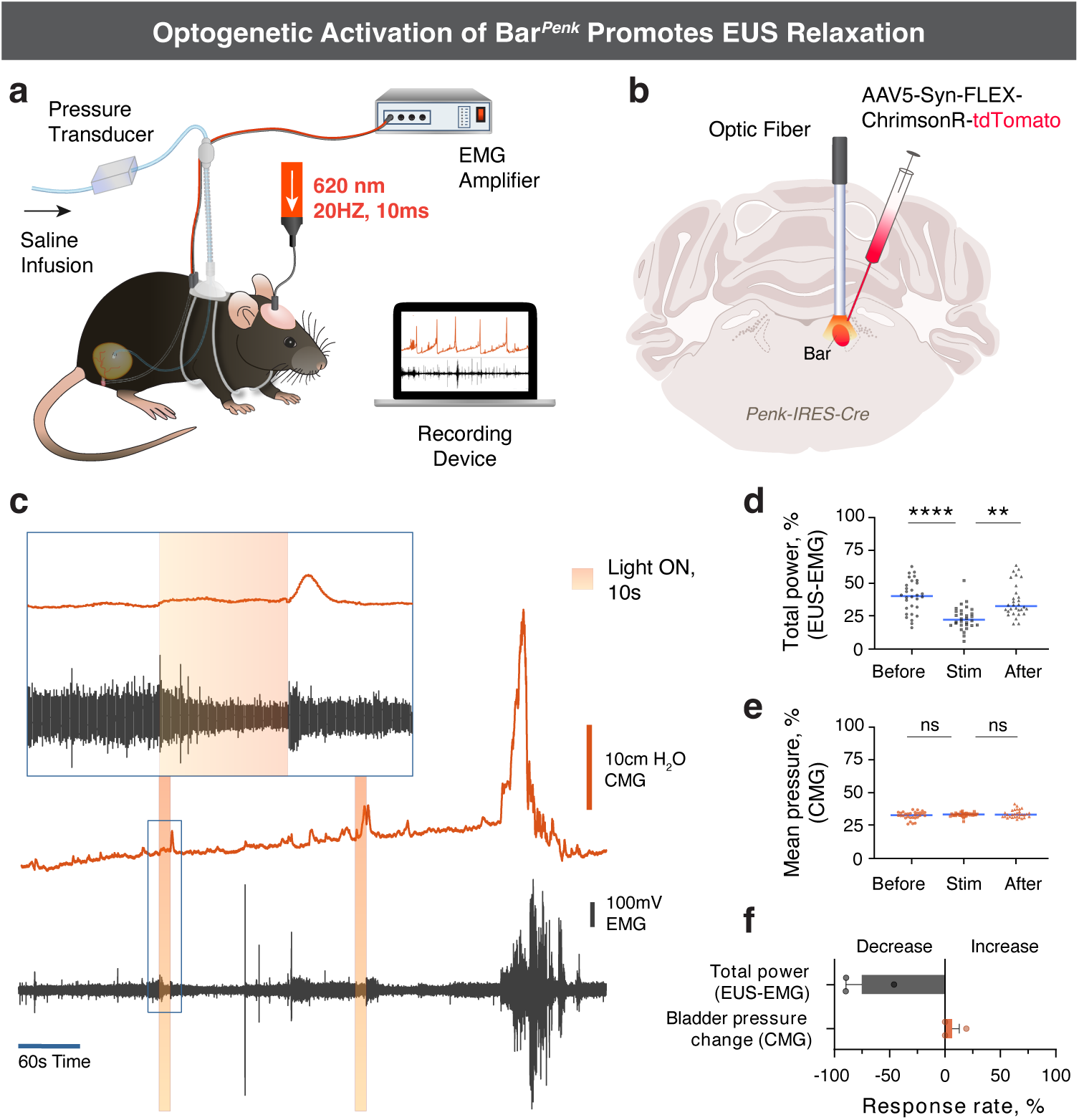
Photostimulation of Bar*^Penk^* neurons promotes relaxation of the external urethral sphincter. **a.** Experimental setup integrating optogenetics, cystometry (CMG), and electromyography (EMG to record bladder pressure and EUS activity during Bar*^Penk^* photostimulation. **b.** Schematic of unilateral FLEX-ChrimsonR injection into Bar with an optic fiber above. **c**. Representative urodynamic tracings showing bladder pressure (red) and EUS-EMG activity (black) with 10s light pulses (orange shading), and a magnified 30s window capturing baseline, stimulation, and recovery phases. **d, e.** EUS-EMG total power (**d**) and average bladder pressure (**e**) before, during, and after stimulation (*n = 30 events from 3 mice, equally weighted;* in (**d**), *\*\*\*\*p <0.0001, ***p = 0.0037;* in (**e**), *p = 0.91, p > 0.99; Friedman test followed by Dunn’s multiple comparisons test).* Blue lines indicate medians. **f.** Likelihood of EUS and bladder responses during photostimulation (*mean ± SEM, n = 3 mice, 74 events*). **Abbreviations**: Bar, Barrington’s nucleus; EUS, external urethral sphincter. *See also Supplementary Fig. 7. Source data are provided as a Source Data file*.

### Retrograde tracing reveals the neural circuits underlying voluntary EUS control

Given that Bar*^Penk^*neurons regulate EUS activity and are essential for voluntary scent marking, they may function as descending command neurons that coordinate voluntary EUS control upon integrating excitatory and inhibitory signals from upstream brain regions. To map afferent inputs to *Penk+* neurons, we employed two complementary retrograde tracing strategies using modified rabies virus (RVdG) (**Fig. 8a, b)**. First, we traced upstream inputs to the entire Bar*^Penk^* population (“All *Penk*+”) by injecting helper AAV (FLEX-TVA-mCherry-oG) and, subsequently, RVdG directly into Bar of *Penk-IRES2-Cre* mice. Notable input sites to Bar*^Penk^* neurons included the subcoeruleus nuclei; the entire extent of the ventrolateral PAG (vlPAG; Bregma −4.3 to −5.1mm), lateral PAG (lPAG; Bregma −4.0 to −4.9), and the dorsomedial PAG (dmPAG); the lateral hypothalamic area (LHA, Bregma −1.2 to −2.5); the bed nucleus of the stria terminalis (BST), with GFP+ cells concentrated in the laterodorsal division; and the central amygdala (CeA) (**Fig. 8c-i; Supplementary Table 4, Supplementary Movie**). Interestingly, a small number of helper-negative, GFP+ neurons were also detected in both the contralateral and the ipsilateral Bar, indicating reciprocal connections between the two Bar nuclei, as well as local connectivity within the nucleus. As a second approach, we labeled upstream neurons that selectively connect with the spinally-projecting Bar*^Penk^* neurons by injecting a retrograde helper virus (AAVrg-DIO-TVA-oG-mCherry) into the lumbosacral spinal cord (**Supplementary Fig. 8a**), followed by RVdG injection into Bar (**Fig. 8b**). The input sites corresponded to those identified with the first approach, though fewer GFP+ neurons were detected in the BST and CeA. While this may suggest that not all Bar*^Penk^* neurons send axons to the spinal cord, differences could also reflect technical factors, such as the use of different helper viruses or local transduction versus retrograde viral uptake at the spinal terminals. Furthermore, dense labeling was observed in the midbrain (vlPAG and lPAG) and pons (PPN) and in a defined ventral region of the zona incerta (SubI nucleus), indicating strong innervation of the spinally projecting Bar*^Penk^* neurons (**Fig. 8i; Supplementary Table 4**).

**Figure 8.**
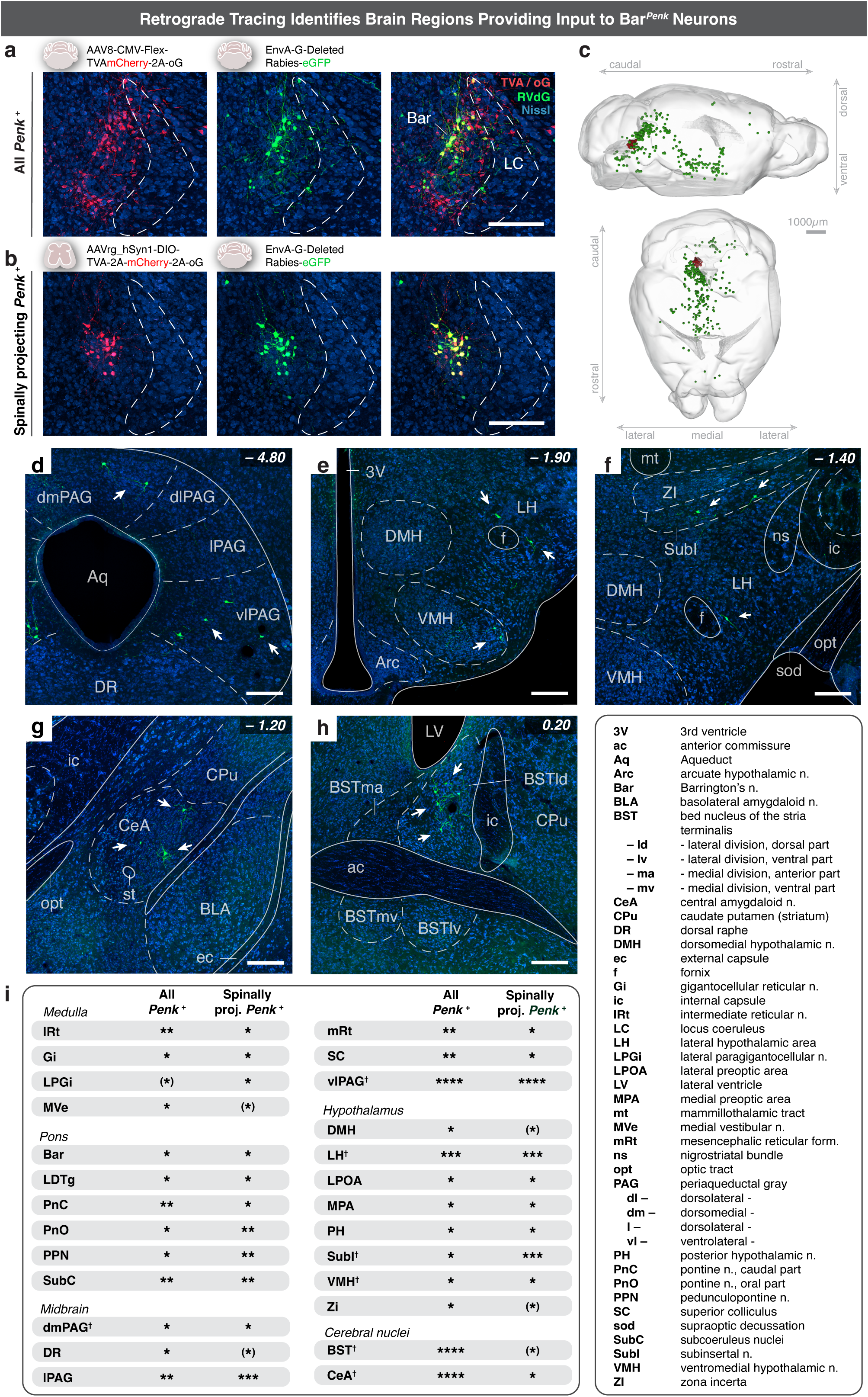
Monosynaptic retrograde tracing of Bar*^Penk^* inputs using modified rabies virus. **a.** Injection of a Cre-dependent helper AAV encoding TVA-mCherry and optimized rabies glycoprotein (oG) into Bar of a *Penk-IRES-Cre* mouse (red, left), followed four weeks later by modified rabies virus (RVdG) injection into the same site (green, middle; *n = 3*). Co-labeled starter cells appear yellow (right). **b.** TVA/oG expression (red, left) in Bar of a *Penk-IRES-Cre* mouse following lumbosacral spinal injection of retrograde AAV encoding DIO-TVA/oG/mCherry, selectively labeling spinally projecting Bar*^Penk^*neurons. RVdG was injected into Bar (green, middle) four weeks later (*n = 4*). Co-labeled starter cells appear yellow (right). **c.** 3D reconstruction of a cleared (iDISCO+) whole brain, displaying Bar*^Penk^* starter cells (red) and RVdG-labeled upstream neurons (green). **d-h.** Representative coronal sections showing RVdG-labeled neurons (green) in key upstream regions: vlPAG (**d**), LH (**e, f**), SubI (**f**), CeA (**g**), and BST (**h**). Approximate Bregma levels are indicated (*n = 7*). **i.** Summary table showing distribution of presynaptic neurons to Bar*^Penk^*, on an arbitrary density scale from 1 to 4 stars. Bracketed stars indicate input regions observed in only a subset of animals. Regions marked with † correspond to panels (**d-h**). In (**a, b, d-h**), neuronal cell bodies are counterstained with Nissl (blue). **Scale bars**: 200 μm, unless noted. **Abbreviations:** form., formation; n., nucleus; proj., projecting. *See also Supplementary Fig. 8, Supplementary Table 4 and Supplementary Movie. Source data are provided as a Source Data file*.

## Discussion

This study advances our understanding of Barrington’s nucleus by providing a multi-scale analysis that defines the cell-type architecture, circuit connectivity, and behavioral roles of its neuronal populations. Our analysis identified five distinct excitatory subtypes, including the well-studied *Crh*-expressing neurons [24, 27, 30], and showed that *Esr1* does not mark a distinct neuronal population in Bar, as previously suggested [25]. We evaluated the specific contributions of the neuronal populations to LUT function; activation of the glutamatergic *Penk*-expressing population revealed striking sex-dependent effects on micturition behavior. Bar*^Penk^* neurons control EUS relaxation in a highly specific manner and are essential for voluntary, socially motivated urinary behaviors such as scent-marking. These findings provide valuable insights into the brainstem circuits that control reflexive and voluntary micturition and highlight how discrete molecular subtypes within Bar contribute to sexually dimorphic regulation of the LUT.

Voiding is initiated when descending output from Bar excites parasympathetic preganglionic neurons in the lumbosacral spinal cord, while simultaneously engaging inhibitory circuits that suppress somatic EUS motor neuron activity. Strong projections targeting the IML and DGC regions have previously been reported for Bar neurons, including Bar*^Crh^* [25–27, 39]. Here we show that four of the identified glutamatergic populations project to the lumbosacral spinal cord: *Crh*, *Penk*, *Tac1*, and *Fgf10*-expressing neurons. Chemogenetic activation of two subtypes - the Bar*^Crh^* and Bar*^Penk^* neurons - increases voiding frequency, consistent with a role in regulating LUT function. The *Tac1+* and *Fgf10+* populations may play modulatory roles or instead contribute to other autonomic functions, for example, colorectal innervation or pain modulation. The glutamatergic *Penk* neurons in Bar, comprising approximately 20% of Bar’s neuronal pool and showing negligible overlap with Bar*^Crh^* neurons, represent a distinct molecular subtype that we characterize in detail here.

In awake and freely behaving animals, Bar neurons are active at specific times during the fill-void cycle [27, 40]. We have previously shown that the intrinsic activity of Bar*^Vglut2^* neurons precedes and then follows the increase in bladder pressure, suggesting that their activation drives micturition [27]. Neural activity in the glutamatergic *Crh* subpopulation in Bar closely tracks bladder pressure during voiding contractions, likely acting to augment detrusor contraction [27]. In contrast, Bar*^Penk^* neurons are active selectively during voiding: their activity rises after the onset of bladder contraction and the duration of this activity closely tracks void duration, suggesting that these neurons are engaged in the ongoing execution of voiding rather than its initiation alone. During non-voiding contractions, which are not accompanied by urine release, Bar*^Penk^* activity is markedly attenuated and typically ceases prematurely. These findings suggest that the neurons may operate downstream of a void-specific gating mechanism, possibly coordinating EUS relaxation only when conditions for micturition are met, and their activity remains inhibited when voiding is contextually inappropriate. Indeed, photostimulation of Bar*^Penk^* neurons suppresses EUS activity in awake mice, confirming that these neurons can reliably induce EUS relaxation.

Among the major projection targets of Bar*^Penk^* neurons is the DGC, which is preferentially innervated relative to the IML. The DGC contains GABAergic/glycinergic interneurons [18, 21] that, when electrically stimulated, evoke relaxation of the EUS [20]. *Penk* neurons likely exert control over the somatic EUS motor neurons, at least in part, via these inhibitory sacral DGC interneurons. During normal voiding, relaxation of the EUS is accompanied by phasic bursting activity, which is thought to facilitate efficient elimination of urine [41, 42]. Our findings suggest that Bar*^Penk^*neurons selectively engage the sacral inhibitory circuit to promote EUS relaxation, without recruiting the spinal bursting center localized in the L3/L4 spinal cord [43, 44]. This is in line with evidence that EUS bursting originates from intraspinal circuitry, as it persists following complete transection above these levels, which eliminates all supraspinal input [45]. Notably, although both Bar*^Penk^* and Bar*^Crh^* neurons project to the DGC and IML, they differentially engage these spinal circuits, with Bar*^Crh^* linked primarily to detrusor activity [25, 27]. Consistent with recent work [46], Bar*^Crh^* also sparsely innervate the superficial dorsal horn, a sensory region implicated in descending pain modulation, whereas Bar*^Penk^* boutons appear more broadly distributed within the dorsal horn region. These differences across functionally distinct spinal domains further highlight the functional heterogeneity of neuronal populations within Bar and underline the need to resolve the molecular identity and organization of their spinal projection targets and local circuitry. The apparent selectivity of Bar*^Penk^* neurons for the EUS relaxation pathway positions them as a promising entry point for dissecting and modulating specific aspects of LUT control in relevant disease models.

Chemogenetic activation of Bar*^Penk^* neurons in male mice produced an aberrant phenotype characterized by numerous leaks, while the frequency of typical, corner-associated voids remained unchanged, albeit with reduced volume per void. Although bladder fullness may influence the extent of leakage, the overall pattern was not consistent with an overflow-incontinence phenotype and instead supports poorly timed, involuntary urine release. Together, these findings suggest that prolonged Bar*^Penk^* activation disrupts bladder-outlet coordination, with frequent and mistimed sphincter relaxation contributing to leakage, while altered coordination between bladder contraction and EUS relaxation may hinder efficient voiding, resulting in smaller void volumes. This interpretation is supported by our optogenetic CMG/EMG recordings, which show that Bar*^Penk^* activation suppresses EUS activity without increasing bladder pressure, indicating that the behavioral effects of chemogenetic stimulation arise from disrupted bladder-outlet coordination rather than direct bladder activation. Our finding that ablation of Bar*^Penk^* neurons had no impact on void frequency, volume, or the ability to maintain continence indicates that *Penk* neurons contribute to, but are not essential for, typical voiding behavior. In fact, ablation of Bar*^Crh^* neurons also does not abolish voiding, despite their proposed role in driving detrusor contractions [24, 25, 27, 30]. Together, these findings may suggest the presence of functional redundancy within Bar. Interestingly, male mice lacking Bar*^Penk^* neurons lose the ability to scent-mark in response to female mouse urine cues, demonstrating their necessity for socially motivated urinary output. Because the detrusor is comprised of smooth muscle cells and is under autonomic control, volitional LUT regulation likely targets the striated muscle of the EUS, which must be actively relaxed to permit urine release [47]. Taken together, these findings position Bar*^Penk^*neurons as a specialized glutamatergic population that coordinates EUS activity both during regular voiding and context-dependent voluntary behaviors.

A key rationale for including both male and female mice in this study stems from the well-established anatomical differences between the male and female LUT system. This sex-specificity is clinically relevant, as LUT conditions, such as overactive bladder and ensuing incontinence, affect aging men and women differently [48], and disorders like Fowler’s Syndrome - a form of urinary retention caused by a poorly relaxing EUS - primarily affects young women [49, 50]. Research directly addressing differences in the neural control of LUT function between sexes remains scarce, but will be essential for advancing clinically relevant insights.

Stimulation of Bar*^Penk^* neurons produces a distinct micturition phenotype in males without an equivalent outcome in females. This raises the possibility that Bar*^Penk^* neurons possess sex-specific characteristics, whether in their molecular identity or circuit connectivity. However, our data does not directly support this interpretation: transcriptomic clustering showed equal contributions from male and female samples, with no evidence of sex-specific subclusters. The proportion of Bar*^Penk^* neurons that are positive for *Vglut2*, or negative for *Crh*, is comparable between sexes, as is the number of *Esr1*-expressing Penk cells. Even though potential differences in hormone sensitivity related to sex-specific variation in circulating or brain estradiol levels cannot be ruled out [51, 52]. Furthermore, axonal tracing revealed similar projection patterns from Bar*^Penk^* neurons in males and females (n = 4M, 4F), and comparable synaptic distribution in lumbosacral regions (n = 4M, 3F), including both the IML and DGC. Kawatani and colleagues analyzed Bar*^Crh^* and Bar*^Esr1^* neuron connectivity with spinal neurons in both sexes, but did not report on sex-specific differences [39].

The observed sex-specific functional output could also arise at the level of downstream circuits. Differences in molecular identity, receptor composition, or excitability of the postsynaptic neurons could influence how the Bar*^Penk^* output is integrated into the spinal circuits. For example, sex-specific variation in opioid receptor expression could result in divergent responses to enkephalin release. Peripheral anatomical differences may also contribute: the EUS is smaller in female mice, and the urethra is shorter [53]. These organs may therefore receive different motor innervation. Lastly, motor neuron pools in females are smaller [54], which may affect the responsiveness of these neurons to descending signals and ultimately influence muscle activation. Future studies are needed to identify sex-specific neuroanatomic and neurophysiological features in LUT signaling, and to determine whether spinal targets of Bar*^Penk^* neurons differ molecularly between sexes.

Both mice and humans maintain voluntary control over urination, and in many mammals, urinary behaviors also serve as a means of social communication via pheromones in urine [22, 23]. Our findings suggest that Bar*^Penk^* neurons facilitate voluntary, scent-marking behavior. Bar receives convergent inputs from multiple brain regions [24, 27, 55], which enables the integration of visceral signals, such as bladder fullness, with contextual cues, to accomplish specific motivated behaviors. The network of upstream sites monosynaptically connecting to Bar*^Penk^*neurons includes regions involved in visceromotor regulation and behavioral state integration, namely vlPAG and LHA [56]. Additional inputs arise from VMH, a sexually dimorphic nucleus implicated in mating behavior [57], MPOA, an area known for its role in social behaviors and previously shown to modulate Bar neurons during socially motivated urination [24], and ZI, which is linked to sensorimotor integration and novelty seeking [58]. Voiding behaviors in general, including scent-marking, require coordinated bladder contraction and EUS relaxation, with different Bar populations serving complementary roles within the voluntary micturition circuit. Consistent with this, the major input sites to Bar*^Penk^* neurons largely overlap with those previously reported for Bar*^Crh^* neurons [24, 27], suggesting that both populations are recruited by shared circuits rather than controlled by distinct upstream nuclei. Their functional divergence may therefore emerge primarily downstream, through differential engagement of spinal targets. Within this broader framework, Bar*^Penk^* neurons may serve as a key node for integrating internal and external cues to coordinate outlet control during context-dependent urinary behaviors.

The vlPAG region is a well-established source of direct projections to Bar [16, 27]. Modulation of this PAG-Bar axis can alter micturition behavior bidirectionally, with glutamatergic signals promoting voiding, and GABAergic ones inhibiting it, consistent with the idea that the PAG serves as a central gate for Bar activity [27, 59, 60]. When voiding is contextually inappropriate, Bar activity remains suppressed despite increasing bladder afferent input to PAG [27, 61]. The sharp rise in Bar*^Penk^* neuron activity at void onset supports a model in which upstream inhibition is transiently lifted to allow sphincter relaxation in synchrony with bladder contraction, facilitating efficient voiding. While PAG likely is an important contributor to this inhibitory gating, additional inputs to Bar [24, 27, 62] may also shape Bar*^Penk^* activity, extending a control network beyond the PAG.

In addition to identifying distinct and functional subpopulations within Bar, our findings refine Bar’s anatomical and molecular boundaries and underscore the need for transcriptomic and spatial resolution when assigning neuronal populations to specific nuclei. For example, while *Esr1* and *Prlr* are expressed across multiple Bar subtypes - including Bar*^Crh^* and Bar*^Penk^* neurons - and their stimulation can reliably promote micturition, neither gene defines a unique population within Bar [33, 46]. Our data indicate that *Esr1* expression is more widespread in and around Bar than previously reported, with substantial overlap between *Esr1* and *Crh*-expressing neurons. A recent study further supports *Esr1* expression across multiple subtypes, revealing distinct Bar*^Esr1^* subgroups: one projecting via the pelvic nerve to regulate bladder-urethra coordination, and another via the pudendal nerve to influence EUS function [32]. Penk, in contrast, marks a molecularly and functionally distinct population with a specific role in LUT control. The previously observed effects on EUS activity with Bar*^Esr1^* stimulation are likely mediated, at least in part, by the *Esr1*+ Bar*^Penk^* neurons.

Limitations of this study include the reliance on DroNc-seq data, which, due to its relatively low sensitivity, may have failed to detect low-abundance transcripts contributing to sex-specific differences in gene expression or missed rare, small populations or subtypes within larger transcriptional clusters. Identification of highly specific molecular markers for Bar may also be inherently limited by the technical constraints of the DroNc-seq approach and by the close anatomical proximity and transcriptional similarity of neighboring pontine populations. Additionally, our tracing studies focused primarily on projections to the spinal cord; future work is needed to map local brainstem circuits and brain-wide afferents to each of the Bar subpopulations, including potential sexual dimorphism. Finally, functional studies under varying physiological and hormonal conditions will be necessary to capture more nuanced differences in LUT regulation between males and females.

Together, these findings establish a cellular and functional framework for understanding how distinct neuronal populations in Bar coordinate autonomic and somatic components of LUT control. This work highlights the intricate brain-body interactions underlying bladder function and identifies Bar*^Penk^* neurons as a specialized population that controls sphincter activity and enables socially motivated micturition in a sex-dependent manner.

## Methods

### Mice

Mice used in this study were bred on a mixed background primarily composed of C57Bl/6J (The Jackson Laboratory). Except for two in-house-generated mouse lines, all mice were obtained from either the Jackson Laboratory or the Lowell Lab (BIDMC, Boston). *Prlr-P2A-Cre* and *Fgf10-IRES-Cre* knock-in mice were generated at the Beth Israel Deaconess Medical Center Transgenic Core Facility, as described below. All transgenic mice included in the functional experiments were used in a heterozygous state.

We used male and female mice ranging from 6 to 25 weeks old (median age: ∼10.5 weeks), with any behavioral and/or histological assessments starting no earlier than 8 weeks of age. Mice weighed between 20 and 32 g. The exact numbers for each genotype used in specific experiments are detailed below and in the figure legends where applicable (see also *Supplementary Table 2*).

To label axonal projections from different neuronal Bar populations to the spinal cord, we utilized male and female mice from the following lines: *Crh-IRES-Cre* (n = 3) [63], *Penk-IRES2-Cre* (n = 8, Jax# 025112), *Tac1-IRES2-Cre-D* (n = 3, Jax# 021877) [64], *Fgf10-IRES-Cre* (n = 4), *Vglut2-IRES-Cre* (n = 3, Jax# 016963) [65], *Vgat-IRES-Cre* (n = 2, Jax# 028862) [65], *Foxp2-IRES-Cre* (n = 4, Jax# 030541) [66], *Prlr-P2A-Cre* (n = 2), and *Esr1-Cre* (Jax# 017911, n = 3) [67]; some of which were crossed with *R26-lsl-L10-GFP* [63].

For chemogenetic (DREADDs) experiments, we used male *Penk-IRES2-Cre* (n = 6), *Tac1-IRES2-Cre-D* (n = 8), *Fgf10-IRES-Cre* (n = 5), and *Prlr-P2A-Cre* (n = 6) mice, some of which were crossed with *R26-lsl-L10-GFP*. To examine the sex-specific effects of chemogenetic activation of Bar*^Penk^*, we additionally included female *Penk-IRES2-Cre* mice (n = 6). To assess the off-target effects of the DREADDs agonists on LUT function, we used male and female *Penk-IRES2-Cre* mice (n = 4 and 6, respectively).

To label and quantify Bar neurons expressing *Crh* and *Penk*, we used adult male *Crh-IRES-Cre* and *Penk-IRES2-Cre* mice crossed to *H2B-TRAP* reporter [68] mice (Jax# 029789) (n = 3 and 3, respectively), while for quantification of *Esr1*-expressing neurons in Bar, we used *Esr1-Cre* mice crossed to *Sun1.sfGFP* reporter *(Jax# 021039, n = 3) [69].* To quantify the overlap of Bar*^Penk^* neurons with previously known Bar markers, we used male and female *Penk-IRES2-Cre* mice crossed to the *ROSA-lsl-tdTomato* (*Ai9*, Jax# 007909) reporter (n = 8) *[70]*. To study the overlap of Bar*^Crh^* with *Esr1*, we utilized adult male *Crh-IRES-Cre* mice crossed to *H2B-TRAP* reporter mice (n = 3) and *Esr1-Cre* (Jax# 017911, n = 2) to validate the anti-Esr1 antibody labeling results. To visualize and quantify the distribution of Bar*^Crh^* and Bar*^Penk^* synaptic terminals in the lumbosacral spinal cord, we used *Crh-IRES-Cre* (n = 4) and *Penk-IRES2-Cre* mice (n = 4 males).

Fiber photometry experiments utilized *Penk-IRES2-Cre* male mice for both all-Bar*^Penk^* (n = 4) and spinally-projecting-Bar*^Penk^* (n = 6) recordings. For experiments investigating the effects of dtA-mediated ablation of Bar*^Penk^* neurons on normal voiding function, we used male and female *Penk-IRES2-Cre* mice crossed to *R26-lsl-L10-GFP*, with n = 8 and 6 in the experimental group and n = 4 and 3 for sham controls. Scent-marking and ablation experiments were conducted using dominant *Penk-IRES2-Cre; R26-lsl-L10-GFP* mice (n = 6 for experimental group (complete ablation), n = 7 for sham control, and n = 6 for the incomplete ablation group).

For optogenetic experiments, we used male *Penk-IRES2-Cre* mice (n = 3). Retrograde tracing with modified rabies studies was performed using male *Penk-IRES2-Cre* mice (n = 7).

### Mouse husbandry

Mice were housed in ventilated racks within air-sealed rooms maintained at a stable temperature (21–23°C) and humidity (30–40%) under a 12-hour light-dark cycle. Each cage contained corncob bedding and nesting material, with ad libitum access to water and standard rodent chow (Teklad F6 Rodent Diet 8664).

In most cases, mice were group-housed after weaning (≤5 per cage), except in the following cases. For scent-marking experiments, male mice were single-housed for at least one week before behavioral assessment. For fiber photometry with cystometry (CMG) and optogenetics with CMG/electromyography (EMG) experiments, mice were singly housed following bladder catheter and/or EMG electrode implantation surgery to minimize implant damage.

All animal care and experimental procedures were approved in advance by the National Institutes of Health and Beth Israel Deaconess Medical Center Institutional Animal Care and Use Committee (Protocol # 024-2023).

### Use of Published Data for Bar Single-Nucleus Atlas

The Bar single-nucleus atlas was generated using a subset of previously published DroNc-seq data from Nardone et al., 2024 [33]. Specifically, we utilized the portion of the dataset corresponding to the micro-dissected Bar (described in the "Mouse Strains and Brain Dissections" section of Nardone et al., 2024). In brief, *Crh-IRES-Cre* mice [63] were crossed with the *L10-GFP* reporter line [63] to generate *Crh-IRES-Cre::R26-lsl-L10-GFP* mice in which *Crh*-expressing neurons were selectively labeled with *GFP*. The highly specific expression of *Crh* in Bar neurons enabled precise 1 mm biopsy punches centered on the *GFP^+^* Bar. Tissue from 18 males and 21 females, processed in 4 and 5 batches, respectively, was used to generate this dataset using DroNc-seq.

We utilized the published digital gene expression (DGE) matrices [33], following standard DroNc-seq processing pipelines. Details on sequencing and data processing, including demultiplexing (bcl2fastq), alignment to the GRCm38 genome (STAR v2.7.3), and UMI collapsing (Hamming distance ≤1), can be found in Nardone et al., 2024. To construct the Bar single-nucleus atlas, 30 DGE matrices from multiple Bar-centered dissections were aggregated and converted into a single Seurat object, with metadata assigned to each batch before merging. Clustering analysis and downstream computational processing were performed by the BNORC Functional Genomics and Bioinformatics Core at Beth Israel Deaconess Medical Center.

### Clustering Strategy and Quality Control

A Single-Cell Remover of Doublets (Scrublet v0.3.2) [71] pipeline was applied to the Bar-DroNc dataset to identify and remove potential doublets. Genes detected in ≤2 cells were filtered out, and nuclei with a mitochondrial gene expression rate >10% and/or <400 unique gene features were removed, as they likely represented empty droplets or low-quality nuclei. This post-filtered dataset of 61,806 nuclei × 26,530 genes was analyzed using Seurat (v3.1.2 [89]; R version 3.6.2; see the “session_info.txt” file in the data repository for a complete listing of all R packages used, together with versioning information). Data were pre-processed using Seurat’s ‘*SCTransform’* function per-sample, which simultaneously performed batch correction, normalization, scaling, and variable feature detection while also regressing out mitochondrial content and inferred cell cycle scores (S and G2/M phases based on known marker gene expression).

Nuclei were then integrated across all samples to ensure proper alignment across different batches. T-SNE (using t-distributed Stochastic Neighbor Embedding) categorical plots were examined to confirm that all samples contributed proportionally to the identified clusters, minimizing batch effect biases. Principal Component Analysis (PCA) was conducted on the 5000 top variable features, followed by clustering and visualization via the aforementioned tSNE plots using the first 30 principal components. A range of parameters was tested to determine the optimal clustering resolution, including 2000, 3000, 4000 or 5000 variable features, 20, 30, or 40 principal components, and clustering resolutions of 0.6, 0.8, 1.0, and 1.2. Cluster stability across resolutions was assessed using *Cluster tree plots*, and the cumulative proportion of variance within each principal component was examined using *elbow plots* to guide PCA cutoff selection. The final clustering resolution was determined in consultation with a bioinformatician.

Differential expression analysis between clusters was performed to identify marker genes. Cell types were assigned to each cluster based on the expression of specific marker genes. Clusters were excluded if they met either of the following criteria: (1) Clusters were considered low-quality if they exhibited very low gene counts and/or high mitochondrial content, (2) clusters were considered mixed or potential doublets if they contained significant markers from multiple unrelated cell types.

After excluding three low-quality or mixed clusters, clustering was performed in three phases to generate a single-nucleus atlas of glutamatergic Bar neurons: (1) all cell types, (2) GABA/Glut neurons, and (3) glutamatergic neurons. In the first phase, the remaining dataset (60,135 nuclei x 26,553 genes) was clustered as described above using a 400UMI cutoff, 5000 variable genes, a 30 principal component cutoff, and a resolution of 0.6. Each cluster of the “All Cells Atlas” (a. 1a-b), was assigned to one of nine major cell types based on the expression of known marker genes. In the second phase, to achieve better separation of excitatory and inhibitory neurons, all non-neuronal clusters were excluded along with neuronal clusters originating from distinct nuclei outside of Bar. The resulting dataset (36,705 nuclei × 26,090 genes) was then re-clustered using the same approach (data not shown).

In the third phase, to generate the final “Atlas of Glutamatergic Putative Bar Neurons”, all GABAergic clusters were excluded based on their expression of the vesicular GABA transporter *Vgat (Slc32a1)*. This was performed iteratively, removing additional GABAergic clusters as they separated. Mixed clusters expressing *Vglut2* (*Slc17a6*) above 10%, regardless of *Vgat* expression, were retained. To reduce the presence of lower-quality nuclei that likely contain a higher proportion of ambient RNA, the UMI cutoff was increased to 800. The final dataset (13,076 nuclei × 25,629 genes) was re-clustered using 2000 variable genes, a 20 principal component cut-off, and a resolution of 0.6.

### Generation of the Knock-in Mouse Lines

*Prlr-P2A-Cre* and *Fgf10-IRES-Cre* knock-in mice were generated at the Transgenic Core Facility at Beth Israel Deaconess Medical Center using the Easi-CRISPR method [72]. A Cre knock-in cassette was inserted into exon 10 of *Prlr* and exon 3 of *Fgf10* using the CRISPR/Cas9 system (**Supplementary Fig. 2a, b**).

### Pronuclear Injection & sgRNA Design

Male C57Bl/6J mice (>8 weeks old, Jax # 000664) were bred with superovulated females to generate zygotes for pronuclear injection. The single-guide RNAs (sgRNAs) were designed using the Benchling platform (https://www.benchling.com/crispr/), and those with the highest scores were selected. sgRNA synthesis was performed using the CRISPRevolution sgRNA EZ Kit (Synthego). Synthego added an 80-mer SpCas9 scaffold to the 20-nucleotide genome targeting sequence in 5’ to 3’ order (excluding the PAM sequence) to create a single guide RNA (1.5 nmol)). Recombinant Cas9 nuclease (#CP01-50) was obtained from PNA Bio. ssDNA templates for *Prlr-2A-Cre* and *Fgf10-IRES-Cre* knock-in cassettes were synthesized by Genscript.

Silent mutations were introduced near the protospacer adjacent motif (PAM) in *Prlr* and *Fgf10* to reduce homology with the wild-type sequence [73]. For effective cleavage at the target site and minimizing non-specific off-target cleavage events, the DNA sequence and target region for the *Prlr* gene was selected to be at position 10329237. On the negative strand, the guide sequence was GCAUGAAGCACGUAGGAUCC with PAM sequence AGG. The knock-in-cassette for *Prlr* contains 147 bp for the 5’ homology arm, the PAM sequence, the modified sgRNA sequence (5’- G GA<u>C</u> CC<u>A</u> AC<u>C</u> TG<u>T</u> TT<u>T</u> ATG C −3’), linker P2A, Cre, NLS, stop codon and 108 bp for the 3’ homology arm. For the *Fgf10* gene, the selected guide sequence was TCTATGTTTGGATCGTCATG with PAM sequence GGG in exon 3 at position 118789301 on the negative strand. The knock-in-cassette for *Fgf10* contained 130 bp for the 5’ homology arm, the PAM sequence, the modified sgRNA sequence including stop codon (5’- C ATG AC<u>C</u> AT<u>T</u> CA<u>G</u> AC<u>T</u> TA<u>A</u> A −3’), IRES2, NLS, Cre, stop codon and 120 bp for the 3’ homology arm.

### Embryo Microinjection & Transfer

Pronuclear-stage embryos were collected and injected following established methods [74–76] using complexes of sgRNA (crRNA + tracrRNA), Cas9, and ssDNA. In brief, the zygotes were microinjected with the aid of an IX71 inverted microscope. Viable zygotes were immediately transferred to a new drop of the M2 medium and then incubated at 37°C in a 5% CO_2_, until reaching the two-cell stage (∼24h post-injection). C57Bl/6J pseudo-pregnant females were used as recipients for embryo transfer, receiving 15–18 microinjected zygotes per oviduct 0.5 days post coitus (dpc). Recipient mothers delivered pups at approximately 19.5 dpc.

### Founder Generation & Genotyping

Of the 260 (for *Prlr*) and 302 (for *Fgf10*) injected zygotes, *221* and *267* were successfully transferred; this resulted in 50 pups and 10 pups born, respectively. DNA sequencing of F0-generation pups revealed that 14/50 of *Prlr* transgenic mice carried the mutant allele instead of the endogenous allele sequence (3 homozygous, 11 heterozygous). As for *Fgf10* transgenic mice, 2 were homozygous, 3 were heterozygous for *Fgf10*-specific Cre insertion, and 1 was positive for generic Cre but not for *Fgf10*-specific Cre, suggesting random genomic Cre insertion(s).

Targeted alleles from founder mice were sequenced to confirm correct knock-in cassette insertion without disruption of endogenous gene expression or function, using Kapa Biosystems HiFi PCR Kit (Roche 07958838001) and TaKaRa Taq DNA Polymerase (Takara HR001A-200U). See Supplementary Table 3 for primers used for Sanger sequencing.

### Backcrossing & Line Establishment

One sequence-confirmed founder (N) was selected for each of the new transgenic lines and backcrossed to wild-type C57Bl/6J mice (Jax # 000664). N1 heterozygous carriers were selected, and their targeted alleles were sequenced (also) and further backcrossed. Resulting F1 heterozygous transgene pups were genotyped (see Supplementary Table 3 for primers) using Roche Taq DNA Polymerase (Sigma Aldrich 4728866001), dNTPack, 100 Units (Sigma Aldrich 728866001). F2–F8 generations were produced by repeated outcrossing with newly purchased wild-type C57Bl/6J mice (Jax # 000664). With generation F8, stable transgenic lines were established and confirmed via sequencing and genotyping.

### General Surgical Procedures

Mice were anesthetized with isoflurane (4% induction, 1–2% maintenance; Kent Scientific VetFlo™) and positioned on a heating mat in a stereotaxic frame (Harvard Apparatus 75-1810). Ophthalmic ointment (Puralub 211-38) was applied to protect the eyes, and either meloxicam-SR (4 mg/kg, s.c.) or buprenorphine-SR (1.2 mg/kg, s.c.) was administered subcutaneously at the start of the procedure, along with antibiotics (Enrofloxacin, 5 mg/kg, s.c.). A sterile field was maintained throughout the surgery, and all surgical tools were sterilized between animals.

After surgery, mice were placed in a clean cage on a heating pad and monitored until fully awake before being returned to their regular housing. Postoperative monitoring was conducted daily, and mice were given at least 21 days for recovery and viral expression before behavioral testing or histological assessment.

For some of the earlier pilot experiments, stereotaxic surgery was performed under ketamine/xylazine anesthesia (100/10 mg/kg, i.p.) using a Kopf stereotaxic frame (David Kopf Instruments 940).

### Viral Injections and Fiber Optic Implantation

Viral vectors were delivered using a Microinjection Syringe Pump System (World Precision Instruments UMP3T-1) with a 10µl Hamilton syringe (Model 1701 RN, Hamilton 7653-01) fitted with a 34G needle (Hamilton 207434).

For some of the earlier pilot experiments, intracranial injections were performed using pulled glass pipettes with an inner diameter of ∼20 µm. An air pressure system was used to deliver picoliter air puffs through a solenoid valve (Clippard EV 24VDC) controlled by a Grass stimulator (Model S48).

### Barrington’s Nucleus Injections

The skin overlaying the skull was shaved and sterilized using sterile alcohol prep pads (Fisher 22-363-750) and povidone-iodide prep pads (Professional Disposables International B40600). A midline incision was made to expose the skull, and after the Bregma-Lambda alignment, a small burr hole was drilled above the target region using a microdrill (Braintree Scientific MD1200120V). The dura was carefully punctured with a pulled glass capillary pipette, and the Hamilton syringe was lowered into the target region. Using stereotaxic coordinates, injections were targeted to Bar at AP: –5.50 mm, ML: ±0.67 mm, DV: –3.85 mm from Bregma. The needle was left in place for 2 minutes before injecting 15–200 nL of the viral vector per side, at a rate of 50 nL/min (exact volumes for specific experiments are detailed below). The needle was left in place for an additional 5–10 minutes before slow retraction. After injections, the incision edges were re-apposed and secured using tissue adhesive (Vetbond 361931).

### Spinal Cord Injections

The surgical area was shaved and sterilized with alcohol and povidone-iodide prep pads. A 10–15 mm incision was made over the T12 to L2 vertebrae, and the L1 vertebra was identified and secured using spinal clamps attached to the stereotaxic apparatus. The connective tissue and muscles were separated from the L1 vertebra to expose the L6–S1 spinal cord by removing the L1 spinous process. The dura was punctured using a 30G needle, and the 10-µl Hamilton syringe fitted with a 34G needle was positioned 0.3 mm lateral to the dorsal spinal vein and lowered 0.5–0.6 mm into the spinal cord. The needle remained in place for 2 minutes before injecting 75 nL at a rate of 50 nL/min. After injection, the needle was left in place for an additional 5–10 minutes before being slowly retracted. A total of six injections were performed, three per side, at different rostral-caudal coordinates across the L6–S1 spinal levels. Following injections, the muscle layer was sutured using 6-0 absorbable vicryl sutures (Ethicon J212H), and the skin was closed using 5-0 non-absorbable polypropylene sutures (Oasis MV-8661).

### Optical Fiber Implantation

Optical fibers were implanted during the same surgery as AAV injections. A Ferrule Holder (RWD Life Science 68214) mounted onto the stereotaxic arm was used to hold and guide a fiber optic cannula along the needle track, as described above, positioning it 0.2-0.3 mm above the injection site. For fiber photometry experiments, stainless steel ferrules (Precision Fiber Products MM-FER2007-304-4500-P) with 400-µm core optical fibers (Thorlabs FT400UMT Multimode, NA 0.39) were implanted unilaterally over Bar. For optogenetics experiments, fiber optic cannulae with Ø1.25 mm ceramic ferrule, 200-µm core (RWD Life Science R-FOC-L200C-50NA) were implanted unilaterally over Bar.

### Viral Vectors: Injection Volumes and Laterality

For anterograde axonal tracing studies, the majority of mice (≥ 2 for each mouse line representing the Bar population) received unilateral injections of AAV8-hSyn-DIO-mCherry (Addgene; plasmid #50459, 15-40nL). Additionally, for some of the pilot cases included here, mice were injected with AAV9-CAG-ChR2(H134R)-mCherry (UPenn; Addgene plasmid #100054), AAV1-Syn-Flex-GCaMP6s-WPRE-SV40 (UPenn; Addgene plasmid #100845), or AAV8-hSyn-DIO-hM3Dq(Gq)-mCherry (Addgene; plasmid #44361). As part of the anti-Esr1 labelling validation, we injected AAV9-hSyn1-DIO-eGFP-2A-FLAG-TeTxLC-WPRE-SV40p (Zurich VVF; v322-9, ∼50 nL). For analysis of synaptic terminal distribution in the lumbosacral spinal cord, mice received a unilateral Bar injection of AAV9-EF1α-DIO-mSyp1-tdTomato-WPRE-bGHp (Zurich VVF; v991-9, ∼30-50 nL).

For fiber photometry recordings from all Bar*^Penk^* neurons, AAV1-Syn-Flex-GCaMP6s.WPRE.SV40 (Addgene; plasmid #100845, ∼30nL) was injected into Bar unilaterally. For recordings specifically from spinally projecting Bar*^Penk^*neurons, mice received 2-4 unilateral spinal injections of the AAVretro-Syn-FLEX-jGCaMP8s-WPRE [77] (Addgene; plasmid #162377, ∼150nL). For chemogenetic experiments, AAV8-hSyn-DIO-hM3D(Gq)-mCherry (Addgene; plasmid #44361, ∼40nL) was injected bilaterally for all mouse lines. For selective activation of spinally projecting Bar*^Penk^* neurons, mice received three bilateral spinal injections of the AAVretro-pEF1a-DIO-FLPo-WPRE-hGHpA [78] (Addgene; plasmid #87306, ∼150nL), followed by bilateral Bar injection of AAV8-hSyn-fDIO-hM3D(Gq)-mCherry-WPREpA (Addgene; plasmid #154868, 100-150nL) 3-4 weeks later. For conditional ablation experiments, AAV8-EF1-lox-Cherry-lox-(dtA)-lox2 (P. Fuller, M. Lazerus; Beth Israel Deaconess Medical Center, ∼40nL) was injected bilaterally, while sham controls received bilateral injections of AAV8-hSyn-DIO-mCherry (∼40nL).

AAV5-Syn-FLEX-ChrimsonR-tdTomato-WPRE-bGHp [79] (Addgene; plasmid #62723, ∼30nL) was injected unilaterally for optogenetic experiments. For monosynaptic conditional retrograde tracing from Bar*^Penk^* neurons, mice received a unilateral injection of the helper vector AAV8-CMV-FLEX-TVAmCherry-2A-oG (Salk; Addgene plasmid #102985, 20–30nL), followed by pseudotyped G-deleted rabies SAD. DG-EnvA-GFP [80] (Salk, Addgene plasmid #32635, 150– 200nL) 3–4 weeks later. For retrograde tracing from spinally projecting Bar*^Penk^* neurons, mice received three bilateral spinal injections of the helper vector AAVretro-hSyn1-DIO-TVA-2A-mCherry-2A-oG-WPRE-bGHp (Zurich VVF; v306-retro, ∼75nL), followed by bilateral Bar injection of SAD-DG-EnvA-GFP (150–200nL) 3–4 weeks later.

### Micturition Video Thermography (MVT)

MVT was performed as described previously [27, 81]. In brief, up to four bottomless, open-top enclosures were placed on filter paper (Cosmos Blotting paper, 360 gsm; Legion Paper P05-COS4060) for each experiment, with an overhead A65 thermal camera (FLIR) capturing the void spot signature. Enclosures were cleaned with Clidox between trials (5-minute contact time, then towel- and air-dried) and high-temperature washed between cohorts to prevent cross-contamination.

Thermal recordings were processed using ResearchIR software, converting sequence files to WMV with embedded timestamps. Voiding events were identified as the first frame of void hotspot appearance, with a screenshot captured once the void spot stabilized (typically 10–15 minutes post-voiding). ImageJ (NIH) was used for volume estimation, with screenshots converted to 8-bit grayscale and thresholded. A 10 cm² reflecting template in each recording calibrated pixel-to-cm² conversion, accounting for camera height and angle variations. Final volume estimates were derived using established calibration curves [81].

To standardize bladder filling, mice received a subcutaneous injection of 1 mL prewarmed 5% dextrose water before recordings and were briefly returned to their home cage (5–10 minutes) for absorption and to prevent filter paper contamination. Mice received a single chow pellet to minimize exploratory behavior during the recording. Water was withheld during recordings to reduce variability in bladder filling and prevent spillage from interfering with void spot analysis. Experiments were conducted in same-sex cohorts of up to four mice, with each placed in its own enclosure. Recordings lasted 2 hours, except for scent-marking assessments (1 hour), as pilot experiments showed dominant mice marked extensively early in the session before subsiding. To control for circadian influences, all recordings were performed at the same time of day, spanning the late light to early dark cycle.

### Typical Mouse Voiding Behavior

A typical voiding sequence begins with the mouse leaving its “home” corner, where it spends most of the test session. The mouse briefly explores, rears, paces, and/or grooms before walking to a corner or wall, turning around to face away from the corner, lifting its tail, and remaining still for several seconds. A fresh urine hotspot then appears on the filter paper, after which the mouse walks away. In control mice, these behavioral features are observed in nearly all voids. Voiding events that deviate from this pattern, such as urine leakage mid-stride, in the center of the cage, or while the mouse is engaged in other behaviors, were categorized as abnormal urine spots (leaks).

In rare cases (<5% of mice, though not formally quantified), we observed animals that appeared unable to exhibit typical conscious voiding behavior and instead showed features of incontinence before any experimental intervention. These mice exhibited (frequent) small leaks during movement, rest, or other behaviors, failed to produce larger voids, or showed signs suggestive of stress incontinence. If these abnormalities persisted across two recording sessions, the mice were excluded from analysis and further experiments.

### Cystometry Bladder Catheter Implantation

Mice were anesthetized with continuous isoflurane (4% induction, 1.5% maintenance). A 1 cm midline abdominal incision was made using a scalpel, and the underlying abdominal muscle was blunt-dissected to expose the bladder. A 3Fr bladder catheter (Instech C30PU-RJV1307) was inserted into the bladder dome and secured with a purse-string suture (8-0 Prolene, DemeTech PM6980065G0P). The catheter was then tunneled subcutaneously to the back of the neck, exteriorized, and attached to a vascular infusion harness (22ga, Instech C30PU-RJV1307). The abdominal muscles were sutured with 6-0 monoacryl sutures (Ethicon Y492G), and the skin was closed with 5-0 non-absorbable sutures (Oasis MV-8661).

Buprenorphine-SR (1.2 mg/kg, s.c.) was administered subcutaneously at the start of the procedure, along with antibiotics (Enrofloxacin, 5 mg/kg, s.c.). After surgery, animals were placed on a heated mat for recovery before being returned to the animal care facility. Mice were monitored and received Enrofloxacin (5 mg/kg, s.c.) daily, until the end of the experiment. To minimize the risk of catheter damage, they were singly housed for the duration of the study. Animals were allowed to recover for 7–10 days before experiments began.

### EUS-EMG Electrode Implantation

If electromyography (EMG) recording was required, electrode implantation was performed during the same surgical procedure. PFA-coated stainless steel wire (37G, A-M Systems 790600) was stripped at one end and shaped into a hook for insertion. Two electrodes were implanted into the external urethral sphincter (EUS), while a ground electrode was sutured to the abdominal muscle. EMG wires were tunneled subcutaneously to the back of the neck, where they were exteriorized and secured. Postoperative recovery followed the same protocol as the catheter implantation, ensuring optimal healing before the experiment started.

### Urodynamic and EUS-EMG Measurements

Urodynamic measurements were performed at approximately the same time of day during the light phase. Awake mice were placed in custom-made acrylic enclosures with filter paper flooring. The bladder catheter was connected to a syringe pump, with an in-line pressure transducer linked to PowerLab for pressure monitoring. EMG electrodes were connected to an amplifier (x1000, 8 Hz to 2 kHz bandpass filter), and EMG signals were sampled using PowerLab. Saline was continuously infused at a rate of 15-30 µL/min for 2-3 hours. Intravesical pressure and EUS-EMG activity were recorded using LabChart software.

### Chemogenetic experiments and analysis

Following hM3Dq AAV injections, mice were allowed 3–4 weeks for recovery and viral expression before testing. Mice received an i.p. injection of either saline (vehicle) or DREADDs agonist 10 minutes before being placed in the MVT behavioral arena. Each session lasted 2 hours, and every mouse underwent the paradigm twice with saline and twice with the agonist, alternating treatments. Behavioral responses were averaged across the two runs.

For Bar*^Penk^* chemogenetic experiments in males, we used Clozapine N-Oxide (CNO, 1 mg/kg, i.p.). We selected a dose at the lower end of the commonly used range (1–5 mg/kg) [25, 27, 30]. For later chemogenetic experiments, we used Compound 21 (C21, 0.8 mg/kg, i.p.), which has been reported to have better brain penetration and possibly fewer off-target effects than CNO [82, 83]. To determine the optimal C21 dosage, we tested a range of 0.5, 0.75, 1.0, and 2.0 mg/kg. Higher doses (>1 mg/kg) induced mild, non-LUT-related side effects, including body temperature changes and decreased locomotion, which were mitigated by using doses ≤1 mg/kg. Therefore, we used C21 in a concentration strictly below 1 mg/kg for all experiments.

To evaluate potential micturition behavior-related off-target effects of these agonists, we administered CNO or C21 to naïve (no DREADDs/Gq present) male and female Penk-IRES2-Cre mice (n = 4 and 5, respectively). Additionally, to assess the efficacy of different agonists, we conducted MVT trials with saline, CNO, or C21 (two trials per condition, n = 2, **Supplementary Fig. 5a**).

### Fiber Photometry Setup

Fiber photometry was conducted as previously described [27] using the Doric Lenses 1-site, 2-color Fiber Photometry system (FPS_1S2C_405/GFP_400-0.48). Calcium-dependent GCaMP6s fluorescence was excited at 465 nm, with the signal modulated at 515 Hz. A 405 nm reference signal was recorded and modulated at 211 Hz to control for motion artifacts and tissue movement. Both excitation wavelengths were coupled through a Doric FMC5 mini cube, and emission light was collected via the same fiber, filtered through the GFP filter, and focused onto a Newport 2151 femtowatt photoreceiver.

The photoreceiver signal was recorded at 4 kHz using LabChart software (AD Instruments). Data were demodulated using an adapted MATLAB script from Doric, downsized to 0.1 s resolution, and further analyzed with custom MATLAB scripts. Bladder pressure (CMG) and GCaMP recordings were synchronized and recorded simultaneously in separate LabChart channels at 4 kHz.

### Experimental Protocol

AAV was allowed to express for at least six weeks before recordings. One-week post-bladder catheter implantation, mice were placed in a metabolic cage with filter paper flooring and connected to the fiber photometry system via an optical patchcord, while the pinport of the harness was linked to a saline infusion pump. Light intensity at the patchcord tip (465 nm) was 0.1–0.2 mW. A thermal camera (FLIR C2) was positioned below the cage to confirm micturition events by the heat signatures on the filter paper following a void.

Each mouse underwent 1-3 hour recording sessions over multiple days, with at least 11 voiding events included per animal. Trials from all sessions were pooled to calculate mean fluorescence changes within 60 seconds before and 60 seconds after micturition events for each mouse. After the experiments, mice were perfused, and brain sections were examined for fluorophore expression and fiber placement. Mice with insufficient gCaMP6s expression in the Bar region or incorrect fiber placement were excluded from the final analysis.

For fiber photometry recordings from spinally projecting Bar*^Penk^*neurons, recordings were performed in conjunction with micturition video thermography (MVT). The spinal cord-injected AAV was allowed to express for 3-4 weeks before optic fiber implantation over Bar. One to two weeks after fiber implantation, mice were volume-loaded with 1 mL of 5% Dextrose (s.c.) and placed in MVT arenas for 2-3 h recording sessions, repeated across multiple days. GCaMP recordings and thermal video were synchronized, and the first thermal video frame in which a urine spot became visible was defined as void onset.

### Fiber Photometry Analysis

Relative fluorescence changes were calculated as ΔF/F₀ = (F - F₀)/ F₀, where F₀ was obtained by applying a best-fit curve to the entire trace. To correct for motion artifacts, the 405 nm (isosbestic) excited GCaMP fluorescence (ΔF/F₀ (405)) was subtracted from the 465 nm calcium-dependent GCaMP fluorescence (ΔF/F₀ (465)). Heatmaps and graphs were generated using custom MATLAB scripts.

To generate an averaged FP-CMG graph, GCaMP traces were normalized per event to account for variability in fluorescence levels between animals and recording days, caused by differences in fiber placement and gradual GCaMP signal decay. The normalization formula used was: (ΔF/F₀ – baseline(ΔF/F₀)) / baseline(ΔF/F₀) × 100, where baseline ΔF/F₀ was defined as the mean ΔF/F₀ during the 60 seconds preceding the event. To create heatmaps, ΔF/F₀ values were converted to Z-scores using the formula: *Z* = (ΔF/F₀ − mean(ΔF/F₀)) / Std(ΔF/F₀).

Fiber photometry recordings from spinally projecting Bar*^Penk^*neurons were processed using the same approach. For correlation analysis between void duration and the duration of peak-plateau neural activity, void duration was defined as the time (in seconds) between the first thermal video frame in which a urine spot became visible and the frame at which the urine spot stopped spreading and/or the mouse left the latrine corner. Peak-plateau duration was defined as the period of sustained-high GCaMP signal following a steep rise in fluorescence shortly before void onset and ending with the subsequent rapid decline in fluorescence (**Supplementary Fig. 4d**).

### Alignment of Voiding and Non-Voiding Events

To align multiple voiding events, we calculated and plotted the differential bladder pressure change rate (dP/dt) and used its maximum value preceding the contraction, representing the steepest rise in bladder pressure before voiding begins, as the alignment point (0-line).

Voiding contractions were aligned to MAX (dP/dt), marking the moment of peak detrusor activity. Voiding contraction events were included in the analysis if the cystometry trace showed a single contraction peak. Non-voiding contractions (NVCs) were aligned to peak bladder pressure. Only NVCs with a single peak with ≥5 cm H_2_O increase in bladder pressure were included to minimize motion-related pressure artifacts. The graph and heatmap display the GCaMP signal from 60 s before to 60 s after the aligned event for both voiding and NVCs.

For random-shuffle graphs, we included 120s of GCaMP traces surrounding randomly selected time points (using the MATLAB randomizer function ‘Randy’), ensuring a comparable number of events while using the same traces as the zero-lag condition.

### Ablation Experiments and Analysis

For DTA-mediated ablation experiments, MVT-recorded behavioral trials were conducted on sexually naïve group-housed mice before AAV injection (POD -1) to establish baseline voiding patterns. Following this initial screening, mice were injected on POD 0 with either a Cre-dependent DTA virus (experimental) or a DIO-mCherry virus (sham control). Both groups underwent identical experimental paradigms and were tested in same-sex cohorts of up to four mice. Before each session, mice were volume-loaded with 1 mL of prewarmed 5% dextrose water subcutaneously and placed in the MVT behavioral arena. Each session lasted 2 hours, with voiding frequency and volume per void recorded for each mouse, following the MVT analysis methods. Behavioral assessments were conducted weekly post-surgery on POD 7, 14, and 21.

Following the final behavioral session, mice were perfused, and brain sections encompassing the core of Barrington’s nucleus (Bar) and adjacent regions were analyzed histologically. Successful ablation was confirmed by a substantial reduction in GFP+ neurons at the injection site, in Penk-IRES2-Cre::L10 mice. Only cases with sufficient bilateral ablation (≥90–95% loss of GFP+ neurons in Bar) and minimal viral spread beyond Bar borders were included in the final analysis.

### Scent-Motivated Marking Behavior

Sexually naïve, group-housed adult *Penk-IRES2-Cre::L10-GFP* male mice were singly housed for at least one week prior to behavioral testing. Before exposure to female scent, mice were habituated to the behavioral arena and screened for abnormal voiding behaviors (e.g., signs of incontinence; see Typical Voiding Behavior section) using the standard MVT setup.

To establish baseline scent-marking behavior, mice underwent two screening sessions before surgery. Before each session, mice were volume-loaded with 1 mL dextrose water subcutaneously, introduced to the behavioral arena, and allowed to explore for 5-10 minutes before being exposed to the stimulus: 100 μL of pre-warmed female mouse urine, pipetted onto filter paper placed in the center of the cage. Scent-marking behavior was then recorded throughout a 1-hour MVT session.

Mice that failed to initiate marking within the first 5 minutes or produced fewer than 10 urine marks during the session were classified as subordinate and excluded from further analysis. The remaining dominant mice were randomly assigned to either the experimental or sham control group (Supplementary Fig. 6e, f) and received bilateral Bar injections of DIO-dtA (experimental) or DIO-mCherry (sham control). 1-hour MVT recordings were conducted in mixed cohorts of four, composed of experimental and control animals.

### Post-Surgical Testing and Analysis

The mice were reintroduced to the behavioral arena three weeks post-injection (POD 21–25), and presented with the urine stimulus again. Each mouse was tested twice on consecutive days, with MVT recordings lasting 1 hour per session. Testing was conducted in mixed cohorts of up to four mice. Thermography videos were analyzed for urine marking behavior, excluding typical corner voids, and latency to marking was noted. Behavioral responses were averaged across both trials. Researchers were blinded to the experimental group during analysis.

Following the final behavioral assessment, mice were perfused, and brain sections encompassing Bar and adjacent regions were analyzed histologically. Successful ablation was confirmed by a substantial reduction (≥90–95%) of Penk-GFP+ neurons at the injection site, with minimal viral spread beyond Bar borders.

### Female Urine Collection

Urine was collected from adult (8–25 weeks) female mice, group-housed up to five per cage. Urine was collected by holding each mouse over wax paper and massaging the bladder (if the mouse did not void spontaneously), then pipetting the urine from the wax paper into a sterile 2mL tube. Urine was pooled from 4–5 cages over four consecutive days to ensure representation of all estrous cycle stages, and briefly stored at −20°C. After the collection period, samples were thawed on ice, pooled in equal proportions, aliquoted, and refrozen at −20°C for use in behavioral assays.

### Optogenetic Experimental Setup

Optogenetic stimulation was delivered using a single-channel LED driver (Plexon, PlexBright LD-1) controlling a 620-nm PlexBright Table-Top LED module (Plexon), delivering 5-10 mW at the fiber tip. The TTL signal was modulated using an Arduino Uno board (Arduino A000066), programmed via Arduino IDE Software to set the frequency and pulse duration. Optical power was measured using a Thorlabs light meter before and after each session to ensure consistency.

For photostimulation experiments, fiber-implanted mice were briefly anesthetized with 4% isoflurane before being connected to optogenetic patch cables (Plexon OPT/PC-FC-LCF-200/230-HP-1.0L KIT) via a ceramic mating sleeve (included in the same kit). The bladder catheter pin port of the harness was connected to a saline infusion pump, and EUS-EMG electrodes were linked to an amplifier. Bladder pressure (CMG), EUS-EMG, and Arduino output recordings were synchronized and recorded simultaneously in separate LabChart channels at 10 kHz.

### Photostimulation Parameters

During pilot experiments, we tested multiple stimulation frequencies (2, 5, 10, and 20 Hz; **Supplementary Fig. 7a**). Based on our observations, photostimulation was delivered using 10-ms pulses at 20 Hz, for 5 or 10 seconds per stimulation. We found that motion-related artifacts interfered with the EMG signal when recorded in awake, freely behaving mice. To minimize this, stimulations were delivered during quiescent periods, avoiding naturally occurring voids. Each mouse underwent 1–2.5-hour recording sessions across multiple days, with at least 19 stimulation events recorded per animal.

### Optogenetic Experiments Analysis

To compare EUS-EMG activity before, during, and after stimulation, the EMG signal (mV) was converted to Total Power (TTP, V^2^), defined as the sum of power across all frequencies within the epoch, using the built-in LabChart function. Mean TTP values were extracted from LabChart for three time periods: before stimulation (baseline, same duration as stimulation), during stimulation (5s or 10s light delivery), and after stimulation (recovery period, same duration as stimulation). To account for variability in voltage across trials and animals, TTP values were normalized using the following formula: TTP_p_= TTP_p_ / (TTP_before_ + TTP_stim_ + TTP_after_) × 100, where TTP_p_ represents the total power during a specific period (before, during, or after stimulation). Similarly, mean bladder pressure (CMG) values were extracted for the same three time periods and normalized using the same approach. To ensure equal representation across animals, an equal number of randomly selected trials (10 per animal) were included using a randomizer function for both Total Power and CMG analyses. Trials were excluded from analysis if movement or electrical/static artifacts were present during the baseline phase (before stimulation). However, trials with potential artifacts during the stimulation or recovery phase were retained to avoid bias in trial selection.

The response rate was calculated per animal. For EMG, the percentage of trials in which a ≥5% decrease in EUS-EMG activity was observed compared to the baseline period (before stimulation). For CMG, the percentage of trials where bladder pressure increased by ≥3 cm H₂O compared to the baseline period, accounting for the natural filling-related pressure increase.

After all experiments, mice were perfused, and brain sections were examined for viral expression and fiber placement. Mice with insufficient Chrimson expression in the Bar region or incorrect fiber placement were excluded from the final analysis.

### Perfusion, Histology Processing, and Assessment

Mice were anesthetized with chloral hydrate (7% w/v in sterile saline, ∼20mg/mL i.p.) and transcardially perfused with phosphate-buffered saline (PBS) at room temperature, followed by 10% formalin (Fisher Scientific 245-685). Brains, and in some cases, spinal cords, were immediately extracted and postfixed overnight in 10% formalin at 4°C. After 24–48 hours of cryoprotection in 25% sucrose, tissues were sectioned using a freezing microtome (Leica SM2000R) in a 1-in-4 series. The whole brain or rostral brainstem were sectioned coronally, while the spinal cord was cut either coronally or horizontally. In most experiments, sections were cut at 30 or 40 μm, except for *in situ* hybridization and neuronal quantification experiments, where sections were cut at 15–25 μm. Sections were stored in cryoprotectant solution at −20°C or in PBS-azide at 4°C before immunofluorescence labeling.

### RNAScope In-Situ Hybridization

Brain tissue was collected using the same perfusion protocol described above, with strictly controlled post-fixation (≤12 hours) and dehydration (12-15 hours) steps. Coronal sections were cut at 15-20 μm, and stored at −20°C until processing according to the protocol provided within RNAscope® Fluorescent Multiplex Reagent Kit (Advanced Cell Diagnostics 320851) user manual.

In brief, brain sections were mounted onto Superfrost Plus microscope slides (Fisher Scientific 12-550-15), washed 3-5 times with distilled water, pretreated with Protease III for 30 minutes, and hybridized with mixed probes (see below). for 2 hours at 40°C, followed by four subsequent amplification steps, each lasting 15-30 minutes. Each of the pretreatment and hybridization steps was followed by 3×2min washes with RNAscope wash buffer. The slides were then air-dried in a light-protected environment, coverslipped using DAPI Fluoromount-G mounting medium (Southern Biotech 0100-20), and stored at −20°C until imaging, typically within 24-48 hours.

List of RNAscope probes used: Mm-*Bnc2*-C2 (ACD 518521); Mm-*Calcr*-C3 (ACD 494071); Mm-*Cdh6*-C2 (ACD 519541); Mm-*Crh*-C2 (ACD 316091); Mm-*Esr1-*C3 (ACD 478201), Mm-*Fign*-C2 (ACD 871521); Mm-*Fgf10*-C2 (ACD 446371); Mm-*Foxp2*-C1 (ACD 428791); Mm-*Inhba*-C2 (ACD 455871); Mm-*Npas1-*C1 (ACD 468851); Mm-*Oprk1-*C2 (ACD 316111); Mm-*Otof-*C2 (ACD 485671); Mm-*Penk-*C3 and Mm-*Penk-*C4 (ACD 318761); Mm-*Prlr-*C3 (ACD 430791); Mm-*Tac1*-C2 (ACD 410351); Mm-*Tfap2b-*C3 (ACD 536371); Mm-*Tnc*-C2 (ACD 465021); Mm-*Slc32a1-*C3 (ACD 319191); Mm-*Slc17a6-*C3 (ACD 319171); Mm-*Slc17a8-*C1 (ACD 431261); Mm-*Sox6*-C2 (ACD 472061); Mm-*UBC-*C3 (positive control, ACD 320881).

### Immunohistochemistry and Nissl Staining

Immunohistochemistry was performed as previously described. In brief, free-floating sections were blocked in 3% normal donkey serum (NDS, Sigma-Aldrich S30) in 0.25% PBST (PBS + 0.25% Triton X-100) for 1 hour, followed by overnight incubation at room temperature with primary antibodies (one to three per experiment) diluted in 3% NDS, 0.25% PBST. The following dilutions were used: anti-ChAT (goat; Millipore AB144P) at 1:250, anti-Esr1 (rabbit; Millipore 06-935) at 1:4000, anti-DsRed (rabbit; Takara 632496) at 1:2000, anti-GFP (chicken; Thermo Fisher A10262) at 1:2000, anti-FOX3 (NeuN, mouse; BioLegend 834501), and anti-TH (mouse; Millipore MAB318) at 1:1000. After washing, sections were incubated with secondary antibodies diluted in 3% NDS, 0.25% PBST for 1–3 hours at room temperature. All secondary antibodies, including anti-Goat (donkey; −488, −594 or −647; Thermo Fisher A-11055, A-11058, A-32849), anti-Rabbit (donkey; −488, −594 or −647; Thermo Fisher A-21206, A-21207, A-32795), anti-Chicken (−488 or −647; Jackson ImmunoResearch 703-546-155, 703-606-155), anti-Mouse (donkey; −488, −594 or −647; Thermo Fisher A-21202, A-21203, A-31571) were used at 1:1000. Each step was followed by three 2-minute PBS washes. For Nissl staining, NeuroTrace Blue (ThermoFisher N21479) or Deep Red (ThermoFisher N21483) was added to the secondary solution at 1:250; or sections were incubated in 0.25% PBST containing NeuroTrace at 1:250 for 1–2 hours.

Sections were then mounted onto gelatin-subbed microscope slides (SouthernBiotech SLD01-CS), air-dried, coverslipped with DAPI Fluoromount-G mounting medium (Southern Biotech 0100-20), and stored at +4°C until imaging.

### Imaging

Whole-slide fluorescence imaging was performed using 10X or 20X objective on an Olympus BX63 scanning microscope with CellSens software (Olympus). OlyVIA software (Olympus) was used to screen cases and identify regions containing transduced soma in Bar or axonal projections to the spinal cord. Cases with missed injections, incomplete hits, extensive spread beyond Bar borders, or incorrect fiber placement were excluded from the analysis. Representative histological images were acquired using a Leica Stellaris 5 scanning confocal microscope with a 10X air or 20X air objective, using LAS X microscope software. Imaging settings were optimized for each experiment to maximize signal range, and z-stack maximum projections were used for representative images and axonal projections.

### Anatomical quantifications

To quantify the percentage of Bar neurons expressing *Penk, Crh*, and *Esr1* we analyzed four consecutive Bar levels (∼15μm thick, ∼100 μm apart) of *Penk-IRES2-Cre::H2B-TRAP, Crh-IRES-Cre::H2B-TRAP, and Esr1-Cre::Sun1.sfGFP* mice (n = 3 per group), counterstained with Nissl. Bar borders were manually delineated bilaterally in Adobe Photoshop (Adobe) based on the distinct oval-shaped boundaries visualized in the Nissl channel, with the reporter channel turned off to avoid bias. The number of Nissl-positive cells within the defined Bar region, representing the total neuronal count, was quantified manually by placing markers on each identified neuron. Reporter-positive cells (*Penk^+^*, *Crh^+^* or *Esr1^+^*) were then quantified using the same approach. The percentage of reporter-positive neurons was calculated as: the number of reporter-positive cells / total Nissl-positive neurons × 100.

To examine the overlap of Bar*^Penk^* neurons with other known Bar markers, we analyzed tissue from male and female *Penk-IRES2-Cre::tdTomato* mice. Three ∼25 μm sections covering the central, rostral, and caudal Bar were included. For *Penk:Esr1* overlap (n = 4M, 3F), immunohistochemistry was performed using an anti-Esr1 antibody (1:4000, Millipore Sigma 06-935). For *Penk:Vglut2* (n = 3M, 3F) and *Penk:Crh* (n = 3M, 3F) co-localization, RNAscope in situ hybridization was performed using *Vglut2* (Mm-Slc17a6-C3; ACDBio 319171) and *Crh* (Mm-Crh-C2; ACDBio 316091) probes. Quantification was performed manually in Adobe Photoshop using a similar approach as above. First, the total number of Penk:tdTomato-positive cells in Bar was quantified, followed by the number of double-positive cells (expressing both tdTomato and the corresponding marker). The percentage of overlap was calculated as follows: the number of double-positive cells / total Penk:tdTomato-positive cells × 100. Researchers were blinded to the extent possible regarding the specific marker being quantified to minimize bias. To quantify and compare Bar*^Penk^* and Bar*^Crh^* synaptic terminal distribution in the lumbosacral spinal cord, we analyzed three consecutive spinal levels (L6, S1, and S2) from *Penk-IRES2-Cre* mice (n = 4M, 3F) and *Crh-IRES-Cre* (n = 4M) injected unilaterally in Bar with DIO-mSyp1-tdTomato. ChAT and NeuN immunostaining were used to identify spinal cord levels. Sections were imaged at 20X on a confocal microscope (Leica Stellaris 5) and processed in Imaris software (Oxford Instruments). Synaptic terminals were thresholded and converted to spots using the same settings for all images. DGC boundaries were manually delineated based on NeuN staining, with the mSyp1-tdTomato channel hidden during ROI assignment to avoid bias. The IML was defined using a standardized circular ROI centered on ChAT-positive bladder motor neurons and applied consistently across sections and animals. Spot counts within each ROI were quantified automatically. Synapse distribution at each level was calculated as: synaptic spots in the ROI (DGC or IML) / total synaptic spots in DGC + IML × 100. For across-level analyses, spot counts in each ROI at a given level were normalized to the total number of synaptic spots detected in DGC + IML across all analyzed levels (L6 + S1 + S2) for that animal.

### Retrograde Tracing Using Modified Rabies Virus

For tracing input sites to Penk+ neurons in Bar (“all *Penk*+”), we introduced a TVA, a receptor for avian leukosis viruses, and optimized rabies glycoprotein (oG) specifically to *Penk*-positive neurons in Bar. This was achieved by focal unilateral injection of Cre-dependent AAV-Flex-TVA-mCherry-oG vector ("helper AAV") into the Bar region of *Penk-IRES2-Cre* mice. After 3–4 weeks for optimal protein expression, G-deleted rabies virus (RVdG) pseudotyped with the avian sarcoma leukosis virus envelope (EnvA), and encoding eGFP was injected into Bar using the same coordinates. For tracing upstream sites of spinally projecting Bar*^Penk^* neurons (“spinally proj. *Penk*+”), a retrograde Cre-dependent helper AAV retrograde viral vector (AAVretro-DIO-TVA-oG-mCherry) was injected bilaterally into the lumbosacral spinal cord (L6-S2 region). After 3-4 weeks of expression, the same EnvA-G-deleted rabies virus (RVdG) was injected bilaterally into Bar. In both cases, seven days after RVdG injection, mice were perfused as described above. After post-fixation in formalin, whole brains were coronally sectioned at 30μm thickness. Every other section was stained with anti-DsRed, Nissl stain, and DAPI, then mounted and scanned as described above. A subset of brains was cleared and stained using the iDISCO+ protocol. These whole brains were then imaged with a light-sheet microscope, followed by 3D reconstruction as described below.

We confirmed a sufficient number of starter cells in Bar (co-labeled with both *mCherry* and *eGFP*). Cases with very few or no starter cells, as well as those with starter cells located outside of Bar borders, were excluded from further analysis. Putative presynaptic neurons expressing *eGFP* (only) were quantified and manually assigned to specific brain regions based on the Paxinos and Franklin Atlas (4th edition) [84], using landmarks visualized with Nissl stain and/or DAPI, and tissue autofluorescence. The number of neurons within each brain structure was normalized to estimate whole-brain values by applying a correction factor, i.e., multiplying by 2 for coronally sectioned brains, where quantification was performed on 2 out of 4 series.

### iDISCO+ tissue clearing

Seven days after rabies virus injection, mice were perfused as described above. Brains were dissected and post-fixed overnight at 4°C in 10% formalin F, then stored in PBS-azide (PBS + 0.02% sodium azide) at 4°C. All incubation steps were performed in 5 mL tubes filled to the top. Brains were processed according to the iDISCO+ protocol [85]. In brief, samples were dehydrated through a graded methanol / ddH_2_O series (20%, 40%, 60%, 80%, 100%, 100%) at room temperature (RT) for 1 hour per step. They were then chilled in 100% methanol at 4°C for 20 minutes before overnight incubation in 66% dichloromethane / 33% methanol at RT with shaking. Following two washes in 100% methanol, samples were bleached overnight at 4°C in 100% methanol containing 5% H_2_O_2_. Rehydration was performed through a descending methanol / ddH_2_O series (80%, 60%, 40%, 20%) for 1 hour per step, followed by a 1-hour PBS wash.

Two additional washes were performed in 0.2% PBST (PBS + 0.2% Triton X-100) before incubation in a permeabilization solution (40 mL PBS, 80 µL Triton X-100, 1.15g glycine, 10mL DMSO, and 0.02% sodium azide, for a total of 50 mL stock solution) for 2 days at 37°C with shaking. Samples were then incubated in a blocking solution (42 mL PBS, 84 uL Tween-20, 0.42mg Heparin, 3 mL normal donkey serum, 5 mL DMSO, and 0.02% sodium azide, for a total of 50 mL stock solution) for 2 days at 37°C with shaking.

For immunolabeling, samples were incubated for 7 days at 37°C with shaking in antibody solution (46 mL PBS, 92 µL Tween-20, 0.46 mg Heparin, 2.5 mL DMSO, 1.5 mL Normal Donkey Serum (NDS), and 0.02% sodium azide) containing primary antibodies (Chicken anti-GFP (1:1000), Rabbit anti-DsRed (1:1000)). This was followed by washing in PBS-based washing solution (50 mL PBS, 0.1 mL Tween-20, 0.5 mg Heparin, and 0.02% sodium azide, for a total of 50 mL stock solution), with five solution changes over 5 hours. Secondary antibody incubation (antibody solution + Donkey anti-Chicken AlexaFluor 647 + Donkey anti-Rabbit AlexaFluor 594) was performed for 7 days at 37°C with shaking, followed by a washing step in the same washing solution with five solution changes over 5 hours, plus an overnight wash.

Samples were then cleared by dehydration through an ascending methanol / ddH_2_O series (20%, 40%, 60%, 80%, 100%, 100%) at RT for 1 hour per step, followed by a 3-hour incubation in 66% dichloromethane / 33% methanol with shaking at RT. Residual methanol was removed by two 30-minute washes in 100% dichloromethane or until samples fully sank. Finally, samples were immersed in dibenzyl ether (DBE) for refractive index matching.

### Lightsheet imaging and 3D reconstruction

Cleared whole-mount mouse brains were imaged using a LaVision BioTec Ultramicroscope II. Samples were immersed in a DBE-filled imaging chamber to maintain refractive index matching and imaged using a 2× zoom objective. The voxel resolution was 1.62 µm × 1.63 µm × 5 µm in the x-, y-, and z-axes, respectively. Image stacks (16-bit TIFF format) were stitched and reconstructed in 3D using Imaris software (Oxford Instruments).

### Quantification and statistical analysis

Statistical analyses were performed using Prism 9 (GraphPad, San Diego, CA, USA), with tests selected based on Prism Software recommendations, and in consultation with the Statistical Support Core at Beth Israel Deaconess Medical Center. Nonparametric tests that avoid assumptions about data distributions or variances were used for all experiments. Statistical parameters, including sample size (n), arithmetic mean, standard error of the mean (mean ± SEM), statistical tests, and significance values, are reported in the Figures and Figure Legends. Statistical significance is indicated in figures as follows: *, p < 0.05; **, p < 0.01; ***, p < 0.001; ****, p < 0.0001; ns (not significant). Details on quantification and analysis of behavior, anatomy, fiber photometry, cystometry, and electromyography data are provided in the corresponding sections.

Sample sizes were not predetermined by statistical methods but were chosen based on prior studies [25, 27] and standard practices in animal behavior experiments. All animals were housed under identical conditions, and littermates were randomly assigned to experimental groups. Investigators scoring behavioral recordings were blinded to recording conditions. Animals were excluded from analysis if histological validation showed poor or absent reporter expression or excessive spread, with exclusion criteria established before data processing. Final sample sizes (n) reflect the number of validated animals per group.

## Resource availability

### Lead contact

Requests for further information and resources should be directed to and will be fulfilled by the lead contact, Anne M.J. Verstegen.

### Materials availability

Mouse lines generated in this study will be made available upon request but may require a completed material transfer agreement.

### Data Availability

This paper analyzes publicly available data from the GEO database (accession code <u>GSE226809</u> [33]). Processed snRNA-seq Bar datasets and associated metadata have been deposited on Zenodo and are publicly available at https://doi.org/10.5281/zenodo.15225406 as of the date of publication. Source data are provided with this paper.

### Code Availability

The Seurat pipeline used for analysis has been deposited on Zenodo and is publicly available at https://doi.org/10.5281/zenodo.15225406 as of the date of publication. Custom scripts used for other analysis are provided with this paper.

## Acknowledgements

The authors thank Dr. Linus Tsai, Dr. Christopher Jacobs, Harini Srinivasan, and the Functional Genomics and Bioinformatics Core at Beth Israel Deaconess Medical Center (BIDMC) for their assistance with snRNA-seq analysis; Dr. Da-qing Wang and the BIDMC Transgenic Core Facility for support in generating transgenic mouse lines; Dr. Clif Saper’s Lab for the generous use of shared confocal equipment, as well as BIDMC Confocal Imaging Core and Neurobiology Imaging Facility at Harvard Medical School for imaging resources. We thank the BIDMC Statistical Support Core for guidance with data analysis, and Dr. Mark Zeidel and members of the lab for helpful feedback and support throughout the project.

## Author contributions

Conceptualization: A.M.J.V. and N.K.; Methodology: N.K., A.M.J.V., and A.M.S.; Investigation: N.K., A.M.S., M.L., A.M.J.V., J.M., C.S., and R.L.; Formal analysis: N.K., A.M.S., M.L., A.M.J.V., and C.S.; Software: N.K., A.M.S., and J.M.; Writing – original draft: A.M.J.V., N.K. and C.S.; Writing – Review & Editing: A.M.J.V., N.K., A.M.S., C.S., M.L. and R.L.; Visualization: N.K., A.M.S., R.L. and A.M.J.V.; Data Curation: N.K. and A.M.J.V.; Supervision: N.K. and A.M.J.V.; Project Administration: A.M.J.V.; Funding acquisition: A.M.J.V.

