## Supplementary Figures and Tables for "“Descending Enkephalinergic Neurons in Barrington’s Nucleus Gate Sex-Biased Control of Micturition”"

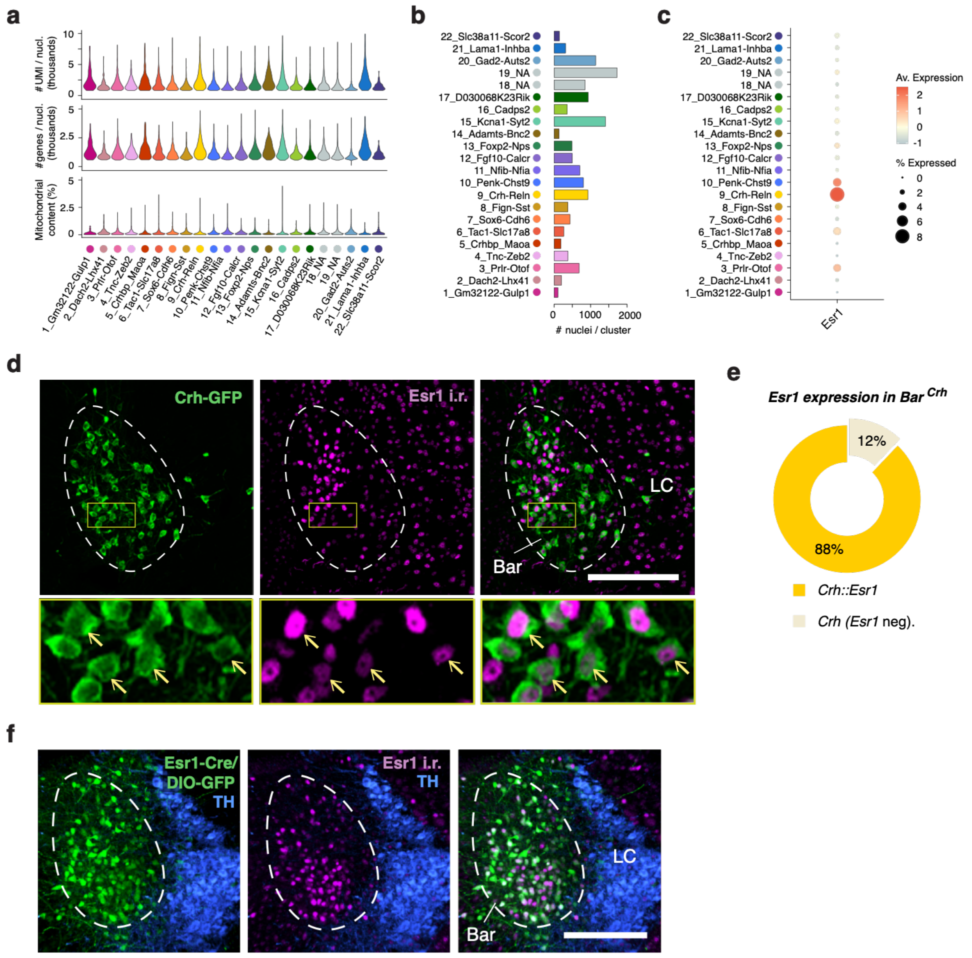

**Supplementary Figure 1. Quality control of snRNA sequencing and *Esr1* expression in Bar.** *Related to Fig. 1.*

**a.** Violin plots displaying quality control metrics for each of the 22 glutamatergic neuronal clusters: number of unique molecular identifiers (top), number of detected genes (middle), and percentage of mitochondrial gene expression (bottom). **b.** Number of nuclei per cluster for the glutamatergic subset of the snRNA-seq dataset. **c.** Dot plot showing *Esr1* expression across the 22 glutamatergic clusters. **d.** Overlap between Crh-GFP (green) and Esr1-immunoreactive neurons (magenta) in Bar of a *Crh-IRES-Cre* mouse crossed to a reporter line; lower panel shows zoomed-in view of the boxed regions above, with yellow arrows indicating neurons co-expressing *Crh* and *Esr1*. **e.** Proportion of *Crh*+ neurons in Bar that co-express Esr1, as detected by immunohistochemistry (*n = 3; 88% ± 1%*). **f.** Expression of DIO-GFP (green) in Bar of an *Esr1-Cre* mouse; Esr1-immunoreactive cells are shown in magenta. Catecholaminergic neurons in the LC are stained for tyrosine hydroxylase (TH, blue).

**Scale bar:** 200 µm. **Abbreviations:** Bar, Barrington's nucleus; i.r., immunoreactive; LC, locus coeruleus; TH, tyrosine hydroxylase; sn-RNA-seq, single-nucleus RNA sequencing; UMI, unique molecular identifier. *Source data are provided as a Source Data file.*

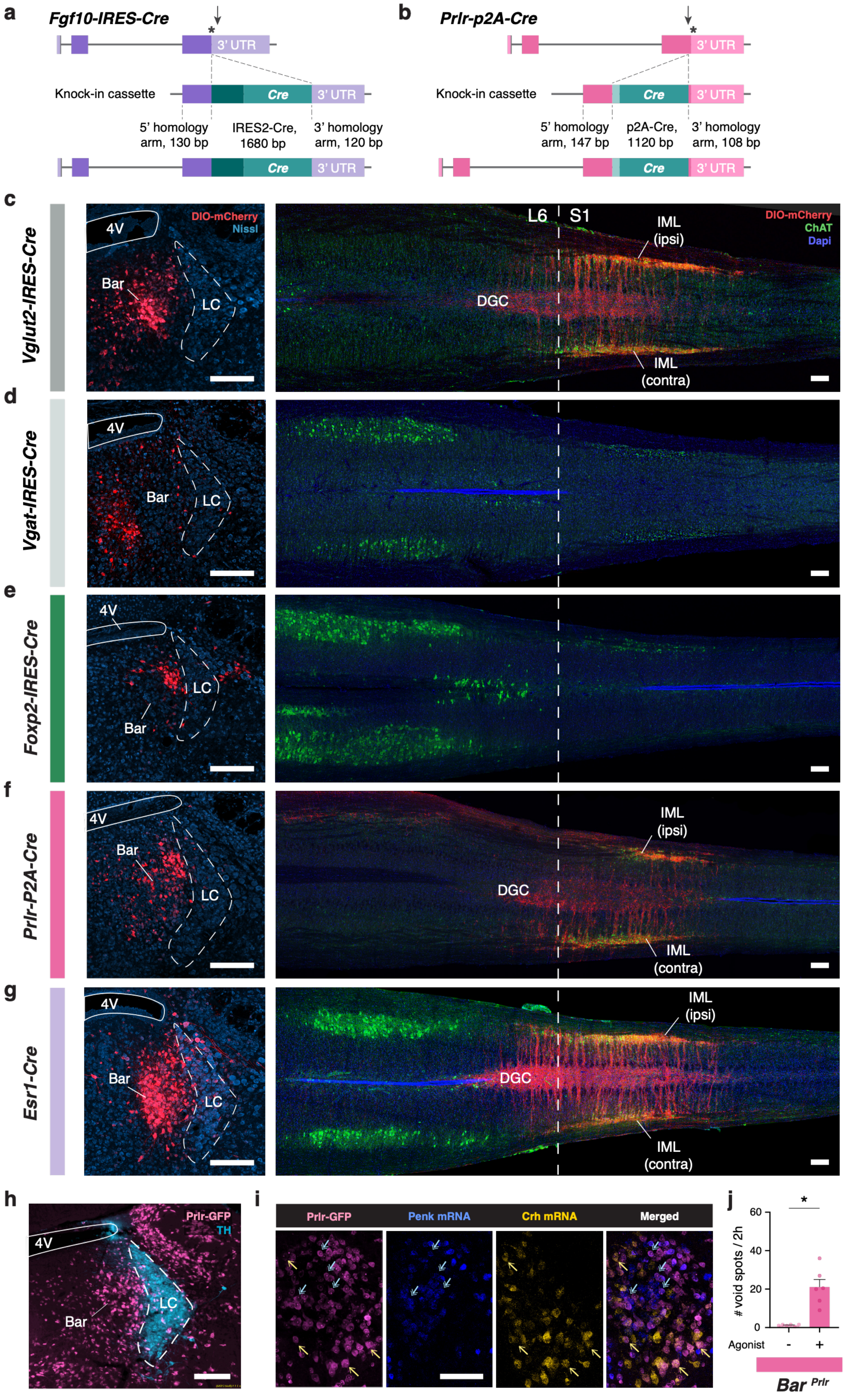
a
Fgf10-IRES-Cre
3' UTR
Knock-in cassette
Cre
3' UTR
5' homology arm, 130 bp
IRES2-Cre, 1680 bp
3' homology arm, 120 bp
Cre
3' UTR
b
Prlr-p2A-Cre
3' UTR
Knock-in cassette
Cre
3' UTR
5' homology arm, 147 bp
p2A-Cre, 1120 bp
3' homology arm, 108 bp
Cre
3' UTR
c
Vglut2-IRES-Cre
4V
DIO-mCherry
Nissl
Bar
LC
L6
S1
IML (ipsi)
DIO-mCherry
ChAT
Dapi
DGC
IML (contra)
d
Vgat-IRES-Cre
4V
Bar
LC
e
Foxp2-IRES-Cre
4V
LC
Bar
f
Prlr-P2A-Cre
4V
Bar
LC
IML (ipsi)
DGC
IML (contra)
g
Esr1-Cre
4V
Bar
LC
IML (ipsi)
DGC
IML (contra)
h
Prlr-GFP
TH
4V
Bar
LC
i
Prlr-GFP
Penk mRNA
Crh mRNA
Merged
j
### void spots / 2h
60
40
20
0
*
Agonist
-
+
Bar Prlr

**Supplementary Figure 2. Characterization of additional putative Bar populations.** *Related to Fig. 2.*

**a, b.** Schematics of knock-in (KI) cassette inserts for *Fgf10-IRES-Cre* (**a**) and *Prlr-P2A-Cre* (**b**) mouse lines generated using the Easi-CRISPR method. The KI cassette for *Prlr* includes a 147 bp 5' homology arm, P2A linker, Cre, NLS, stop codon, and a 108 bp 3' homology arm. The KI cassette for *Fgf10* includes a 130 bp 5' homology arm, a stop codon, IRES2, NLS, Cre, stop codon, and a 120 bp 3' homology arm. **c-g.** Injection of DIO-mCherry (red) in Bar of *Vglut2-IRES-Cre* (**c**; *n = 3*), *Vgat-IRES-Cre* (**d**; *n = 2*), *Foxp2-IRES-Cre* (**e**; *n = 4*), *Prlr-P2A-Cre* (**f**; *n = 2*), and *Esr1-Cre* (**g**; *n = 3*) mouse lines (left panels), with neuronal cell bodies counterstained with Nissl (teal blue). Corresponding axonal projections (or lack thereof) to lumbosacral spinal cord levels are shown (right panels). Ipsilateral (ipsi) and contralateral (contra) projections are indicated when present. Cholinergic neurons are stained for ChAT (green), and nuclei are counterstained with DAPI (blue). **h.** Prlr-GFP expression (pink) in Bar of *Prlr-P2A-Cre* mice crossed with an L10-GFP reporter, showing the distribution of *Prlr*-expressing neurons in and around Bar. Catecholaminergic neurons in the locus coeruleus (LC) are stained for tyrosine hydroxylase (TH, teal blue). **i.** *Penk* mRNA (blue, second panel) and *Crh* mRNA (yellow, third panel) expression in Prlr-GFP+ neurons (pink, first panel) in Bar. The merged image (fourth panel) shows the overlap of *Prlr* expression with other Bar markers. Yellow arrows indicate neurons co-expressing *Prlr* and *Crh*, while blue two-headed arrows highlight *Prlr-Penk* colocalization. **j.** Micturition frequency recorded during 2h MVT trials in *Prlr-IRES-Cre* mice injected with DIO-hM3Dq following saline or hM3Dq agonist administration (*mean ± SEM, n = 6, *p = 0.031, Wilcoxon matched-pairs signed-rank test*).

**Scale bars:** 200 µm. **Abbreviations:** 4V, 4th ventricle; Bar, Barrington's nucleus; bp, base pair; ChAT, choline acetyltransferase; DGC, dorsal gray commissure; IML, intermediolateral column; KI, knock-in; LC, locus coeruleus; MVT, Micturition Video Thermography; NLS, nuclear localization sequence; TH, tyrosine hydroxylase; UTR, Untranslated Region. *See also Supplementary Table 2. Source data are provided as a Source Data file.*

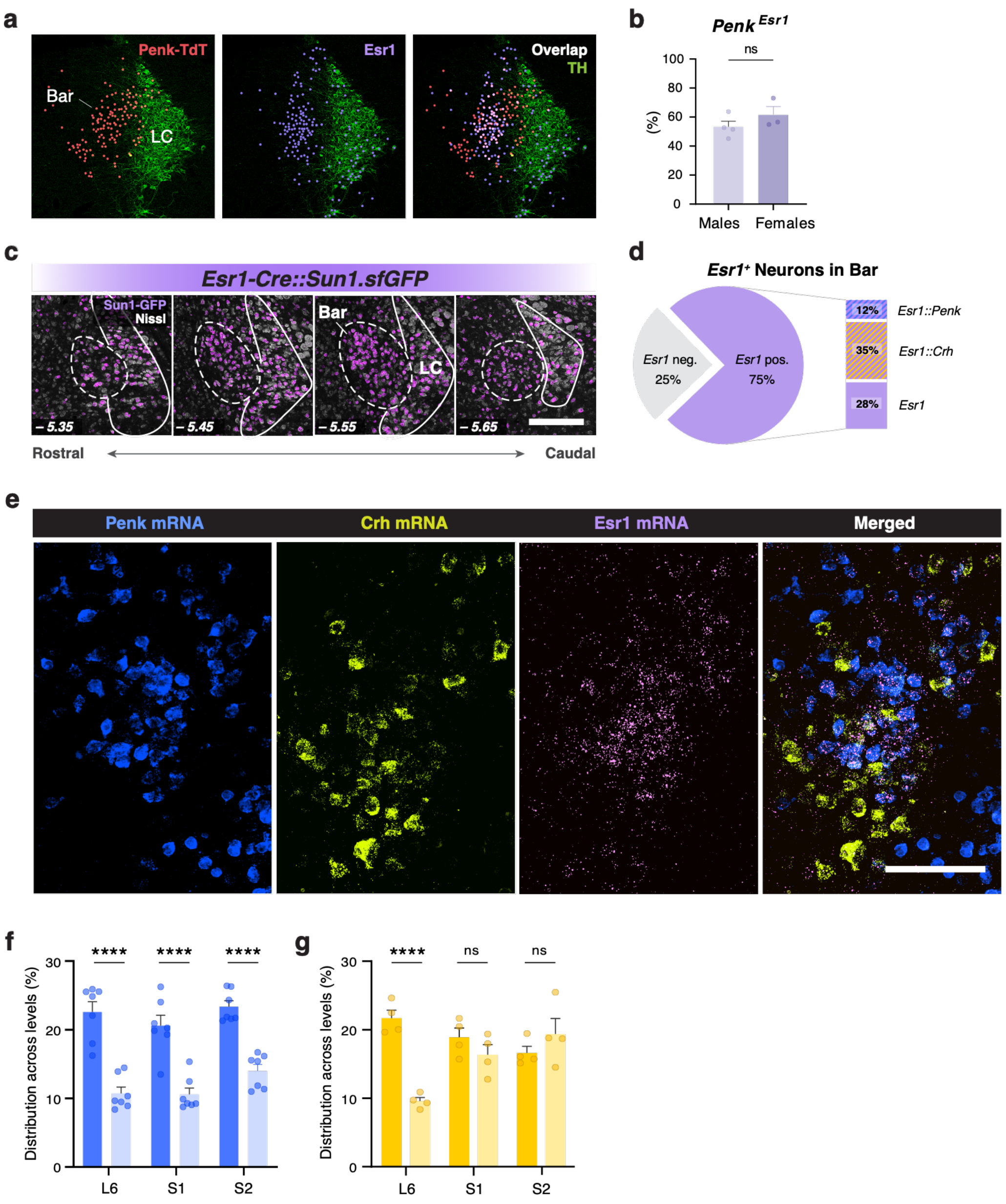


**Supplementary Figure 3. Molecular and anatomical characterization of Bar$^{Penk}$ neurons.** *Related to Fig. 3.*

**a.** Schematic representation of Penk-TdTomato (red, left panel) and Esr1-immunoreactive neurons (magenta, middle) distribution in Bar. The merged image (right panel) shows their overlap (white), with TH-positive neurons of LC (green) included as an anatomical landmark. **b.** Bar graph comparing the percentage of neurons co-expressing *Penk* and *Esr1* in Bar for males (*53.3 ± 3.9%, n = 4*) and females (*61.4 ± 5.8%, n = 3*). *Mean ± SEM, p = 0.23, Mann-Whitney test*. **c.** Distribution of *Esr1+* (magenta) neurons across Bregma levels in Bar; neuronal cell bodies are counterstained with Nissl (gray). **d.** Left, proportion of Esr1-expressing neurons in Bar shown as a pie chart (*75% ± 1.5%, n = 3*). Right, schematic breakdown of the Bar$^{Esr1}$ population inferred from measured overlap with Bar$^{Penk}$ and Bar$^{Crh}$ populations, showing contributions of *Esr1::Penk* (12% of total Bar neurons), *Esr1::Crh* (35% of Bar), and *Esr1* neurons negative for both markers (28% of Bar). **e.** *Penk* mRNA (blue, first panel), *Crh* mRNA (yellow, second panel), and *Esr1* mRNA (pink, third panel) expression in Bar, with merged image shown on the right (scale bar 100μm). **f, g.** Distribution of synaptic terminals in the DGC and IML, expressed as the percentage of total synapses across all analyzed spinal levels for $Bar^{Penk}$ (**f**; *n = 7, ****p < 0.0001 for all*) and $Bar^{Crh}$ (**g**; *n = 4, ****p < 0.0001, p = 0.51, p = 0.44, respectively). Mean ± SEM, one-way ANOVA with Sidak's multiple-comparisons test.*

**Scale bars:** 200 μm, unless noted. **Abbreviations:** Bar, Barrington's nucleus; DGC, dorsal gray commissure; IML, intermediolateral column; LC, locus coeruleus; TdT, TdTomato; TH, tyrosine hydroxylase. *Source data are provided as a Source Data file.*

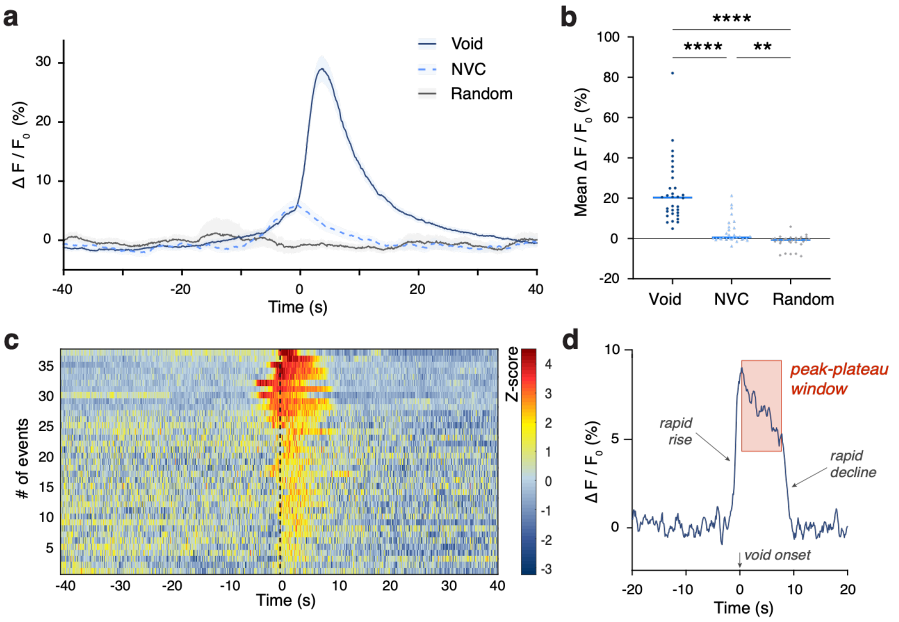


**Supplementary Figure 4. Fiber photometry-based $Ca^{2}$+ imaging of $Bar^{Penk}$ during voiding and non-voiding contractions**. *Related to Fig. 4.*

**a.** Averaged gCaMP6s signals (*$\Delta F/F_0$*) aligned to the onset of voiding (solid blue line), the peak of non-voiding contractions (dashed light blue line), and randomly shuffled time points (solid gray line). Data are presented *as mean ± SEM (n = 86, 73, and 44 events, respectively, from 4 mice), with shading representing the standard error of the mean.* **b.** Mean gCaMP6s signal (*$\Delta F/F_0$*) during voiding events, non-voiding contractions, and random time points. *Thick blue lines indicate the median (n = 32 events from 4 mice, equally weighted; ****p < 0.0001, ****p < 0.0001, **p = 0.0074, Kruskal-Wallis test followed by Dunn's multiple comparisons test)*. **c.** Heatmap of GCaMP8s signal recorded from spinally projecting $Bar^{Penk}$ neurons before, during, and after individual voiding events (*z-scored $\Delta F/F_0$; n = 37 events from 6 mice*), sorted by peak signal. **d.** Representative void-aligned fiber photometry trace highlighting peak-plateau neural activity (red shaded region), defined as the sustained elevated GCaMP signal between the rapid rise and subsequent rapid decline in fluorescence.

**Abbreviations**: NVC, non-voiding contraction. *Source data are provided as a Source Data file.*

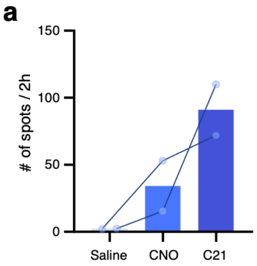


**Supplementary Figure 5. Efficacy of different DREADD agonists**. *Related to Fig. 5.*
**a.** Comparison of the effects of different DREADD agonists on LUT function in mice expressing hM3Dq receptor in Bar$^{Penk}$ neurons. Micturition frequency was recorded during 2-hour MVT trials following administration of saline, CNO, or C21 (*n = 2 mice, two trials per condition (averaged)*).

**Abbreviations:** C21; Compound 21; CNO, clozapine N-oxide; DREADD, Designer Receptors Exclusively Activated by Designer Drugs. *Source data are provided as a Source Data file.*

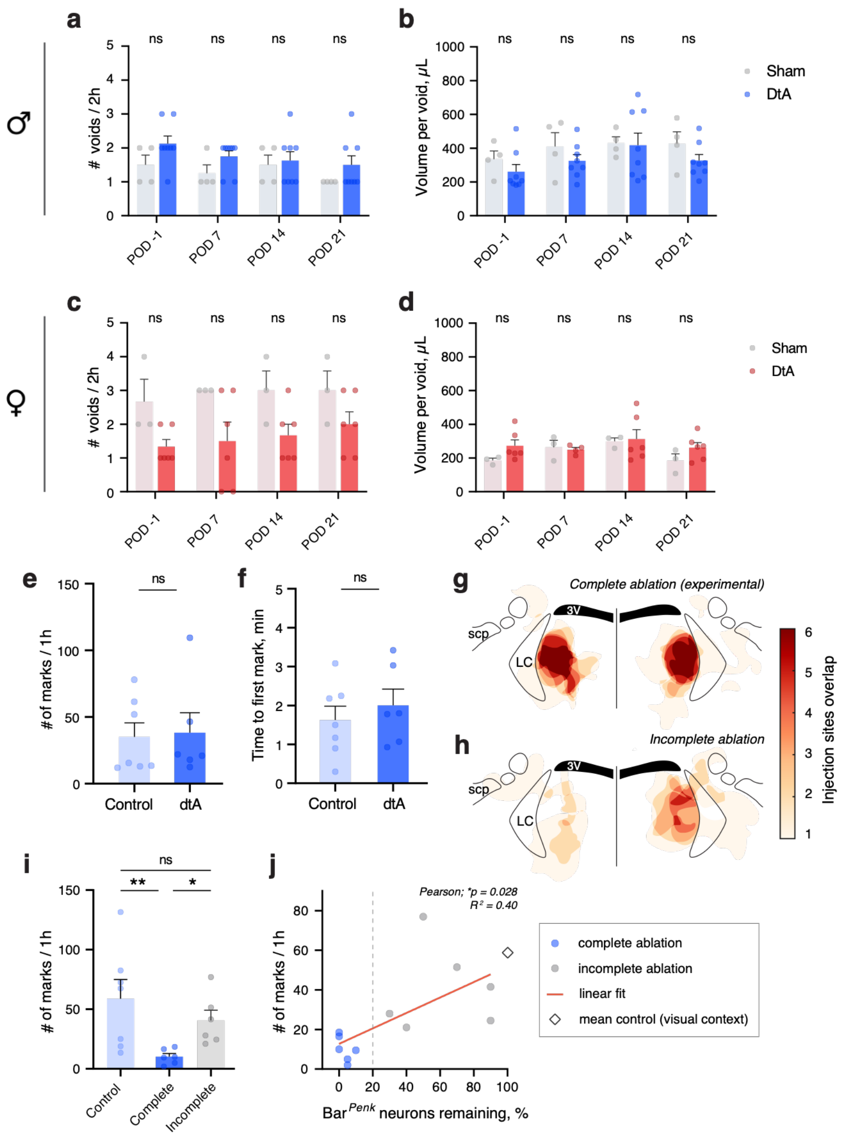


**Supplementary Figure 6. Extended time course of Bar$^{Penk}$ ablation effects and pre-ablation scent-marking behavior**. *Related to Fig. 6.*

**a, c.** Average voiding frequency during 2h MVT trials before and after ablation of Bar$^{Penk}$ neurons compared to sham controls for males (**a**; *n = 8 vs. n 4; p = 0.57, p = 0.57, p > 0.99 p = 0.66*) and females (**c**; *n = 6 vs. n = 3; p = 0.33 for all PODs*). *Mean ± SEM, multiple Mann-Whitney tests with Holm-Šídák's correction for multiple comparisons*. **b, d.** Average urine volume per void recorded MVT trials before and after ablation of Bar$^{Penk}$ neurons compared to sham controls for males (**b**; *n = 8 vs. n 4; p = 0.75, p = 0.74, p = 0.75, p = 0.75*) and females (**d**; *n = 6 vs. n = 3; p = 0.33, p =0.86, p = 0.86, p = 0.60*). *Mean ± SEM, multiple Mann-Whitney tests with Holm-Šídák's correction for multiple comparisons*. **e, f.** Baseline scent-marking behavior: Average number of urine marks **(e)** and latency to the first mark **(f)** recorded during pre-ablation MVT trials (POD -1) before AAV injection of either DIO-dtA (experimental group) or DIO-mCherry (control), (**e, f**; *n = 6 vs. n = 7; p = 0.73, p = 0.63, respectively; mean ± SEM, Mann-Whitney test*). **g, h.** Injection site overlap for the complete ablation cohort (**g;** experimental, *n = 6;*) and incomplete ablation group (**h**; partial, unilateral/mistargeted Bar coverage, *n* = *6*), reconstructed from mCherry-labeled Cre-negative cells within the injection field around the dtA-ablated $Bar^{Penk}$ neurons. **i.** Average number of urine marks during 1h MVT trials on POD 21-25 in control (mCherry; *n* = *7*), complete bilateral $Bar^{Penk}$ ablation (*n* = *6*), and incomplete ablation groups (*n* = *6*). *Mean ± SEM, control vs complete: **p = 0.009; complete vs incomplete: *p = 0.02; control vs incomplete: p >0.99, Kruskal-Wallis, followed by Dunn's multiple comparisons tests.* **j.** Correlation between the percentage of $Bar^{Penk}$ neurons remaining after ablation and marking events. Linear regression was performed across dtA-injected animals only (*n = 12; Pearson r = 0.63;* $R^2 = 0.40$*; *p = 0.028)*; regression line shown in red. Dashed line indicates the predetermined complete-ablation inclusion criterion; control mean (open diamond) is shown for visual context only.

**Abbreviations:** Bar, Barrington's nucleus; dtA, diphtheria toxin A; MVT, Micturition Video Thermography; POD, postoperative day. *Source data are provided as a Source Data file.*

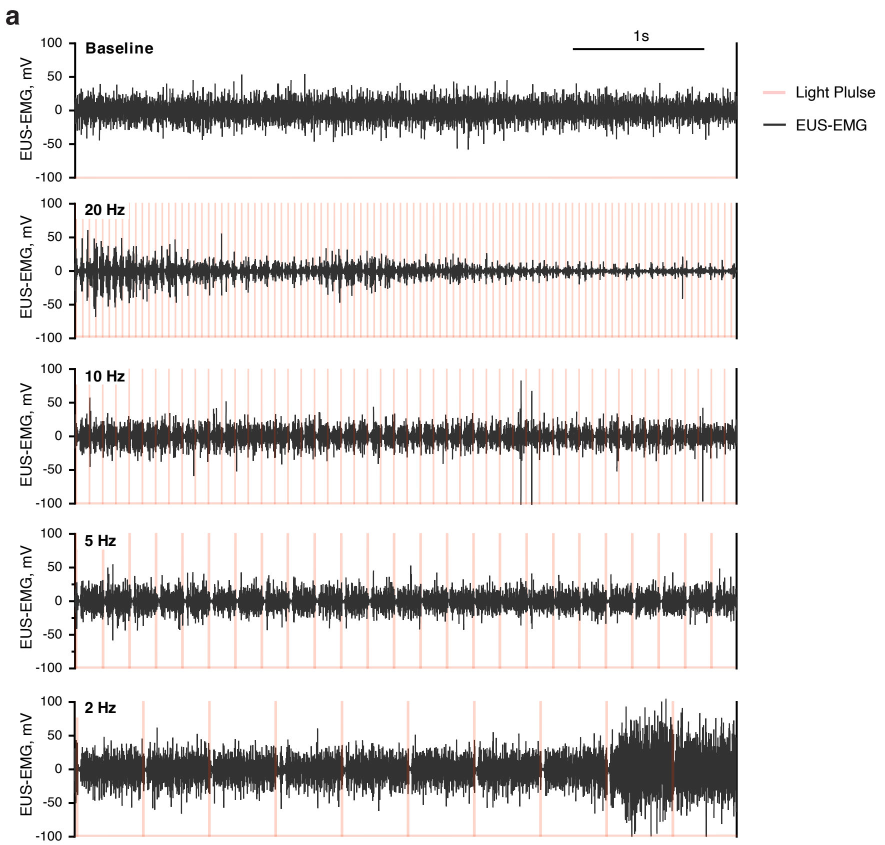


**Supplementary Figure 7. Effects of optogenetic stimulation at different frequencies.** *Related to Fig. 7.*

**a.** EUS-EMG activity recorded in response to different photostimulation frequencies applied to $Bar^{Penk}$ neurons expressing FLEX-ChrimsonR. Transparent red bars indicate light pulses delivered via an Arduino-controlled system.

**Abbreviations:** EUS, External Urethral Sphincter; EMG, Electromyography. *Source data are provided as a Source Data file.*

**a**

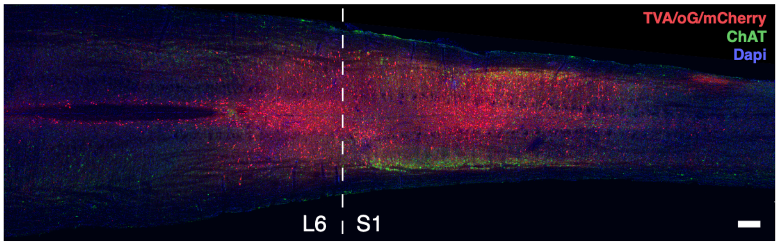


**Supplementary Figure 8. Monosynaptic rabies tracing from spinally projecting Bar$^{Penk}$ neurons.** *Related to Fig. 8.*
**(a)** Representative histological image showing the injection site of a retrograde helper AAV encoding a Cre-dependent construct for TVA (avian receptor for leukosis viruses), optimized rabies glycoprotein (oG), and mCherry (red) in the lumbosacral spinal cord of a *Penk-IRES2-Cre* mouse. Cholinergic neurons are stained with ChAT (green), and nuclei are counterstained with DAPI (blue). The dashed white line indicates the L6/S1 level of the spinal cord.

**Scale bar:** 200 µm. **Abbreviations:** ChAT, choline acetyltransferase.

| Cluster Name | Marker Genes | Source | Summary | Included in Further Analyses? |
|---|---|---|---|---|
| **1_Gm32122-Gulp1** | NA | NA | No Unique or Verifiable Markers (Noncoding RNA or NA) | Excluded |
| **2_Dach2-Lhx4** | *Dach2, Lhx4* | ABA ISH data | Detected outside of Bar | Excluded |
| **3_Prlr_Otof** | *Prlr, Otof* | RNAScope ISH | Detected within Bar / In close proximity | Included |
| **4_Tnc-Zeb2** | *Tnc, Zeb2* | RNAScope ISH, ABA ISH data | Detected outside of Bar | Excluded |
| **5_Crhbp_Maoa** | *Maoa, Th* | IHC, ABA ISH data | Detected outside of Bar | Excluded |
| **6_Tac1-Slc17a8** | *Tac1, Slc17a8* | RNAScope ISH | Detected within Bar / In close proximity | Included |
| **7_Sox6-Cdh6** | *Sox6, Cdh6* | RNAScope ISH | Detected outside of Bar | Excluded |
| **8_Fign-Sst** | *Fign, Sst* | RNAScope ISH, ABA ISH data | Detected outside of Bar | Excluded |
| **9_Crh-Reln** | *Crh, Reln, Npas1* | RNAScope ISH, ABA ISH data | Detected within Bar / In close proximity | Included |
| **10_Penk-Chst9** | *Penk, Tfap2b* | RNAScope ISH | Detected within Bar / In close proximity | Included |
| **11_Nfib-Nfia** | *Nfib, Nfia* | ABA ISH data | Sparse / Widespread Expression | Excluded |
| **12_Fgf10-Calcr** | *Fgf10, Calcr, Oprk1* | RNAScope ISH | Detected within Bar / In close proximity | Included |
| **13_Foxp2-Nps** | *Foxp2, Lhx9* | RNAScope ISH | Detected within Bar / In close proximity | Included |
| **14_Adamts-Bnc2** | *Bnc2* | RNAScope ISH | Detected outside of Bar | Excluded |
| **15_Kcna1-Syt2** | *Kcna1, Syt2* | ABA ISH data | Sparse / Widespread Expression | Excluded |
| **16_Cadps2** | *Cadps2* | ABA ISH data | Detected outside of Bar | Excluded |
| **18_NA** | NA | NA | No Unique or Verifiable Markers (Noncoding RNA or NA) | Excluded |
| **17_D030068K23Rik** | NA | NA | No Unique or Verifiable Markers (Noncoding RNA or NA) | Excluded |
| **19_NA** | NA | NA | No Unique or Verifiable Markers (Noncoding RNA or NA) | Excluded |
| **20_Gad2-Auts2** | NA | NA | No Unique or Verifiable Markers (Noncoding RNA or NA) | Excluded |
| **21_Lama1-Inhba** | *Inhba, Grin2c* | RNAScope ISH, ABA ISH data | Detected outside of Bar | Excluded |
| **22_Slc38a11-Scor2** | *Slc38a11, Dpy19l1* | ABA ISH data | Sparse / Widespread Expression | Excluded |

**Supplementary Table 1. Marker Genes Used for Spatial Mapping of Putative Bar Populations.** *Related to Fig. 1.*
Summary of marker genes used to categorize each of the 22 clusters based on spatial expression patterns, determined using RNA In Situ Hybridization (ISH) or immunohistochemistry (IHC), and/or publicly available ISH image data from the Allen Mouse Brain Atlas (ABA, Allen Institute for Brain Science, 2004). Clusters categorized as "Detected within Bar / In close proximity" were included in further analysis, while those categorized as "Detected outside of Bar" and "Sparse / Widespread Expression," or "No Unique or Verifiable Markers (Noncoding RNA or NA)" were excluded from further consideration. Specific markers for each cluster are provided, and the inclusion or exclusion of a cluster from further analysis is noted.

| Mouse line / Cohort | n (M/F) | Figure panels |
|---|---|---|
| **Quantification of Crh::Esr1 overlap** | | |
| *Crh-IRES-Cre::L10-GFP* | 3 (3/0) | Supp. Fig. 1d, e |
| **Validation of anti-Esr1 antibody staining** | | |
| *Esr1-Cre* | 2 (2/0) | Supp. Fig. 1f |
| **Axonal projection labelling** | | |
| *Crh-IRES-Cre* | 3 (3/0) | 2c |
| *Penk-IRES2-Cre* | 8 (4/4) | 2d |
| *Tac1-IRES2-Cre* | 3 (2/1) | 2e |
| *Fgf10-IRES-Cre* | 4 (4/0) | 2f |
| *Vglut2-IRES-Cre* | 3 (1/2) | Supp. Fig. 2c |
| *Vgat-IRES-Cre* | 2 (2/0) | Supp. Fig. 2d |
| *Foxp2-IRES-Cre* | 4 (4/0) | Supp. Fig. 2e |
| *Prlr-P2A-Cre* | 2 (2/0) | Supp. Fig. 2f |
| *Esr1-Cre* | 3 (2/1) | Supp. Fig. 2g |
| **Functional screening of Bar populations w/ DREADDs** | | |
| *Penk-IRES2-Cre* | 6 (6/0) | 2i (left) |
| *Tac1-IRES2-Cre* | 8 (8/0) | 2i (middle) |
| *Fgf10-IRES-Cre* | 5 (5/0) | 2i (right) |
| *Prlr-P2A-Cre* | 6 (6/0) | Supp. Fig. 2j |
| **Quantification of Crh+, Penk+, Esr1+ neurons in Bar** | | |
| *Crh-IRES-Cre::H2B-TRAP* | 3 (3/0) | 3a, c, e |
| *Penk-IRES2-Cre:: H2B-TRAP* | 3 (3/0) | 3b, c, e |
| *Esr1-Cre::Sun1.sfGFP* | 3 (3/0) | Supp. Fig. 3c, d |
| **Quantification of synaptic terminal distribution** | | |
| *Penk-IRES2-Cre* | 7 (4/3) | 3f - h, Supp. Fig. 3f |
| *Crh-IRES-Cre* | 4 (4/0) | 3f - h, Supp. Fig. 3g |
| **Overlap of Bar$^{Penk}$ neurons with previously known Bar markers** | | |
| *Penk-IRES2-Cre::TdTomato /* Penk::Vglut2 overlap | 6 (3/3) | 3d (left) |
| *Penk-IRES2-Cre::TdTomato /* Penk::Esr1 overlap | 6 (3/3) | 3d (middle); Supp. Fig. 3a, b |
| *Penk-IRES2-Cre::TdTomato /* Penk::Crh overlap | 7 (4/3) | 3d (right) |
| **Fiber photometry recordings** | | |
| *Penk-IRES2-Cre* / All | 4 (4/0) | 4c - f; Supp. Fig. 4a, b |
| *Penk-IRES2-Cre* / Spinally projecting | 6 (6/0) | 4h - k; Supp. Fig. 4a |
| **Chemogenetic activation of Bar$^{Penk}$** | | |
| *Penk-IRES2-Cre* / Experimental males | 6 (6/0) | 5a - e, i |
| *Penk-IRES2-Cre* / Agonist control males | 4 (4/0) | 5c - e |
| *Penk-IRES2-Cre* / Experimental females | 7 (0/7) | 5f - h |
| *Penk-IRES2-Cre* / Agonist control females | 6 (0/6) | 5f - h |
| *Penk-IRES2-Cre* / Experimental / spinally projecting | 7 (7/0) | 5k, l |
| **Effects of Bar$^{Penk}$ ablation on voiding** | | |
| *Penk-IRES2-Cre::L10-GFP* / Experimental (dtA) males | 8 (8/0) | 6b - d; Supp. Fig. 6a, b |
| *Penk-IRES2-Cre::L10-GFP* / Sham (mCherry control) males | 4 (4/0) | 6b - d; Supp. Fig. 6a, b |
| *Penk-IRES2-Cre::L10-GFP* / Experimental (dtA) females | 6 (0/6) | 6e, f; Supp. Fig. 6a, b |
| *Penk-IRES2-Cre::L10-GFP* / Sham (mCherry control) females | 3 (0/3) | 6e, f; Supp. Fig. 6a, b |
| **Scent-marking behavior and Bar$^{Penk}$ ablation** | | |
| *Penk-IRES2-Cre::L10-GFP* / Experimental (dtA, complete) | 6 (6/0) | 6i - k; Supp. Fig. 6e - h |
| *Penk-IRES2-Cre::L10-GFP* / Sham (mCherry control) | 7 (7/0) | 6i - k; Supp. Fig. 6f - h |
| *Penk-IRES2-Cre::L10-GFP* / Incomplete ablation | 6 (6/0) | Supp. Fig. h |
| **Optogenetic stimulation of Bar$^{Penk}$** | | |

| | | |
|---|---|---|
| *Penk-IRES2-Cre* | 3 (3/0) | 7e - f; Supp. Fig. 7a |
| **Retrograde tracing** | | |
| *Penk-IRES2-Cre* / All | 3 (3/0) | 8a, c, g - i; Supplementary Table 4 and 5 (video) |
| *Penk-IRES2-Cre* / Spinally projecting | 4 (4/0) | 8b, d - f, i; Supp. Fig. 8a; Supplementary Table 4. |

**Supplementary Table 2. Summary of mice used in each experiment.** *Related to Online Methods.*
For each experiment, the mouse line/cohort, sample size (n), sex (M/F), and corresponding figure panels are listed.

| Sanger sequencing Knock-In alleles primers | | |
|---|---|---|
| **Prlr_ Primers #** | **Forward_ Sequence (5' -> 3')** | **Reverse_ Sequence (5' -> 3')** |
| Prlr F3 | TGGCCAGCTTTACTGCAACC | |
| Cre R3 | | GCTAGAGCCTGTTTTGCACG |
| Cre F2 | AATGGTTTCCCGCAGAACCT | |
| Cre R6 | | CAGCGTTTTCGTTCTGCCAA |
| Cre F3 | TTGGCAGAACGAAAACGCTG | |
| Prlr R4 | | AGCAGACCACATTACTTCATGACT |
| **Fgf10_ Primers #** | | |
| 11F | TTTAACTGGCAGCACAATGGC | |
| 11R | | GCATCCTTCAGCCCCTTGTT |
| 2F | CAACGTCTGTAGCGACCCTT | |
| 2R | | AGGTTCTGCGGGAAACCATT |
| 3F | AATGCTTCTGTCCGTTTGCC | |
| 3R | | CGGTGCTAACCAGCGTTTTC |
| 4F | TGGCATTTCTGGGGATTGCT | |
| 4R | | ACCATTGCCCCTGTTTCACT |
| 14F | GCGCCCTGGAAGGGATTTTT | |
| 14R | | TCCATCCAAACTTGACTTGCCT |
| **Genotyping primers** | | |
| **Prlr_ Primers #** | **Forward Primer** | **Reverse Primer** |
| Prlr-P2A-Cre (mutant allele); 565bp | (Prlr-F3): TGGCCAGCTTTACTGCAACC | (Cre-R3): GCTAGAGCCTGTTTTGCACG |
| Prlr-WT; 717bp | (Prlr-F3): TGGCCAGCTTTACTGCAACC | (Prlr-R3): TGCATACAGGCAATGGGTCAT |
| Generic Cre; 434bp | (Cre-F2): AATGGTTTCCCGCAGAACCT | (Cre-R5): GGTGCTAACCAGCGTTTTCG |
| **Fgf10_ Primers #** | | |
| Fgf10-IRES-Cre (mutant allele); 1060bp | (Fgf10-F3): GGCAGGCAAATGTATGTGGCATTG | (RVS1 Cre): GCATTGCTGTCACTTGGTCG |
| Fgf10-WT; 618bp | (Fgf10-F3): GGCAGGCAAATGTATGTGGCATTG | (Fgf10-R2): TTGTGCATATGATATGACCCAAGTGC |
| Generic Cre; 434bp | Cre-F2): AATGGTTTCCCGCAGAACCT | (Cre-R5): GGTGCTAACCAGCGTTTTCG |

**Supplementary Table 3. Sanger Sequencing and Genotyping Primers for *Prlr* and *Fgf10* Knock-In Alleles.** *Related to Online Methods.*

List of the primer sequences used for Sanger sequencing and genotyping of *Prlr-P2A-Cre* and *Fgf10-IRES-Cre* knock-in alleles. Separate primer sets are provided for mutant and wild-type allele identification, along with generic Cre `primers used for confirmation.

| Brain Region | Bregma Levels | All *Penk*$^{+}$ | | Spinally projecting *Penk*$^{+}$ | | | |
|---|---|---|---|---|---|---|---|
| | | L7PR | D7PN | G56PR | G57PL | L77PL | L78PRL |
| **Medulla** | | | | | | | |
| **Gi**, gigantocellular reticular nucleus | *-5.7 to -7.2* | – | 4 | 4 | – | 8 | 4 |
| **LPGi**, lateral paragigantocellular nu. | *-5.7 to -7.1* | – | 4 | 2 | 8 | 14 | 2 |
| **MVe**, medial vestibular nucleus | *-5.4 to -6.0* | 4 | 8 | 8 | – | – | – |
| **Rpa**, raphe pallidus | *-6.4; -7.0* | 2 | – | 2 | – | – | – |
| **IRt**, intermediate reticular nucleus | *-5.4 to -6.4; -7.2 to -7.4* | 6 | 20 | 4 | – | 4 | 2 |
| **SN**, Solitary nucleus | *-6.0 to -6.6;* | – | 12 | – | 4 | – | – |
| **Pons** | | | | | | | |
| **CG**, central gray | *-5.3* | – | – | – | – | 2 | 2 |
| **Bar,** Barrington's nucleus (contralateral and interconnected) | *-5.3 to -5.5* | 4 | 4 | 2 | 4 | 4 | 4 |
| **DMTg,** dorsomedial tegmental area | *-4,9; -5.3* | – | – | 2 | – | 2 | – |
| **LDTg,** laterodorsal tegmental nu. | *-5.1 to -5.5* | 6 | – | 4 | 8 | 2 | 2 |
| **LPBn,** lateral parabrachial nucleus | *-5.2* | 2 | – | – | – | – | 2 |
| **MPBn,** medial parabrachial nu. | *-4.9* | – | – | 2 | – | 2 | – |
| **PnC,** pontine reticular nu., caudal | *-4.9 to -5.6* | 12 | 8 | 2 | 8 | 4 | – |
| **PnO,** pontine reticular nu., oral | *-4.3 to -4.8* | 8 | – | 12 | 8 | 8 | 2 |
| **PPN**, pedunculopontine nucleus | *-4.6 to -5.0* | 2 | 4 | 8 | 12 | 8 | 8 |
| **Su5,** supratrigeminal nucleus | *-5.1* | 2 | – | 2 | – | – | – |
| **SubC,** subcoeruleus nuclei | *-5.0 to -5.3* | 4 | 12 | 16 | 8 | 6 | 4 |
| **Midbrain** | | | | | | | |
| **dmPAG,** dorsomedial peri-aqueductal gray | *-3.9 to -4.9* | 2 | – | 6 | 4 | 6 | 2 |
| **lPAG,** lateral periaqueductal gray | *-4.0 to -4.9* | 4 | 4 | 24 | – | 12 | 24 |
| **mRt,** mesencephalic reticular formation | *-3.3 to -3.8; -4.3 to -4.7* | 6 | 16 | 2 | 12 | 6 | 4 |
| **PMnR,** paramedian raphe nucleus | *-4.4 to -4.8* | – | – | 2 | – | – | 4 |
| **SCs,** Superior colliculus | *-3.5 to -3.8; -4.3 to -4.7* | 6 | 12 | 4 | 8 | 4 | 2 |
| **vlPAG,** ventrolateral periaqueductal gray | *-4.3 to -5.1* | 20 | 16 | 64 | 20 | 50 | 44 |
| **Hypothalamus** | | | | | | | |
| **DMH,** dorsomedial hypothalamic nu. | *-2.0 to -2.3* | 8 | – | – | 4 | – | – |
| **LH,** lateral hypothalamic area | *-1.2 to -2.5* | 10 | 16 | 24 | 44 | 10 | 24 |
| **LPOA,** lateral preoptic area | *0.0 to -0.3* | 4 | 4 | 4 | 4 | – | 10 |
| **MPA,** medial preoptic area | *0.1 to -0.2* | – | – | 2 | 8 | 4 | 12 |
| **PH,** posterior hypothalamic nucleus | *-2.1 to -2.5* | 2 | – | 6 | 4 | 2 | 6 |
| **SubI,** subinsertal nucleus | *-1.3 to -1.6* | – | 4 | 18 | 24 | 10 | 14 |
| **VMH,** ventromedial hypothalamic nu. | *-1.7 to -2.0* | 2 | 4 | 2 | 8 | 4 | 12 |
| **Zi,** zona inserta (caudal part) | *-2.5 to -2.8* | 4 | 4 | – | – | 10 | – |
| **Cerebral Nuclei** | | | | | | | |
| **BST,** bed nu. of the stria terminalis | *0.2 to -0.1* | 24 | 28 | 6 | 4 | – | – |
| **CeA,** central amygdaloid nucleus | *-1.2 to -1.7* | 40 | 40 | 10 | – | 4 | 4 |

**Supplementary Table 4. Brain input sites to Bar*$^{Penk}$* neurons.** *Related to Fig. 8.*

Number of rabies-transfected neurons detected across 33 putative input sites to all *Penk*-expressing neurons in Bar (n = 2) and spinally projecting Bar$^{Penk}$ neurons (n = 4). Input sites are categorized by macrostructures, including the medulla, pons, midbrain, hypothalamus, and cerebral nuclei. The second column provides the approximate Bregma span of eGFP-labeled cells.

**Supplementary Movie Legend: "3D Visualization of Upstream Inputs to Bar$^{Penk}$.** *Related to Fig. 8.* A 3D reconstruction of the whole brain, cleared and stained using the iDISCO+ protocol, showing Bar$^{Penk}$ starter cells (red spheres) and RVdG-labeled upstream neurons (green spheres). The video first displays a 360° rotation from a lateral view, followed by a walkthrough in coronal orientation progressing caudal to rostral. Tissue autofluorescence provides structural context. Scale bar shown in lower left corner."
